# Bayesian analysis of GWAS summary data reveals differential signatures of natural selection across human complex traits and functional genomic categories

**DOI:** 10.1101/752527

**Authors:** Jian Zeng, Angli Xue, Longda Jiang, Luke R Lloyd-Jones, Yang Wu, Huanwei Wang, Zhili Zheng, Loic Yengo, Kathryn E Kemper, Michael E Goddard, Naomi R Wray, Peter M Visscher, Jian Yang

**Affiliations:** Institute for Molecular Bioscience, The University of Queensland, Brisbane, Queensland, Australia; Queensland Brain Institute, The University of Queensland, Brisbane, Queensland, Australia; Institute for Advanced Research, Wenzhou Medical University, Wenzhou, Zhejiang 325027, China; Faculty of Veterinary and Agricultural Science, University of Melbourne, Parkville, Victoria, Australia; Biosciences Research Division, Department of Economic Development, Jobs, Transport and Resources, Bundoora, Victoria, Australia

## Abstract

Understanding how natural selection has shaped the genetic architecture of complex traits and diseases is of importance in medical and evolutionary genetics. Bayesian methods have been developed using individual-level data to estimate multiple features of genetic architecture, including signatures of natural selection. Here, we present an enhanced method (SBayesS) that only requires GWAS summary statistics and incorporates functional genomic annotations. We analysed GWAS data with large sample sizes for 155 complex traits and detected pervasive signatures of negative selection with diverse estimates of SNP-based heritability and polygenicity. Projecting these estimates onto a map of genetic architecture obtained from evolutionary simulations revealed relatively strong natural selection on genetic variants associated with cardiorespiratory and cognitive traits and relatively small number of mutational targets for diseases. Averaging across traits, the joint distribution of SNP effect size and MAF varied across functional genomic regions (likely to be a consequence of natural selection), with enrichment in both the number of associated variants and the magnitude of effect sizes in regions such as transcriptional start sites, coding regions and 5’- and 3’-UTRs.

## Introduction

The joint distribution of SNP effect size and minor allele frequency (MAF) is an essential component of the genetic architecture of human complex traits and is influenced by natural selection^1^. A negative relationship between effect size and MAF is a signature of negative (or purifying) selection^2,3^, which prevents mutations with large deleterious effects becoming frequent in the population. Understanding how natural selection has shaped genetic variation helps researchers to improve experimental designs of genetic association studies^4^ and the estimation of SNP-based heritability (the proportion of phenotypic variance explained by the SNPs)^5–8^. Inference on natural selection is also a critical step towards the understanding of the genetic architecture of complex traits. For instance, the theory of negative selection^9^ explains why the effects of common variants identified by genome-wide association studies (GWAS) are unlikely to be large^10,11^.

We have recently developed a Bayesian method (BayesS) to estimate the effect size-MAF relationship, which was considered as a free parameter (*S*) in the model^12^. We detected negative *S* for a number of complex traits in humans, highlighting an important role of negative selection in shaping the genetic architecture, consistent with the findings from other studies based on genome-wide variance estimation approaches^7,10,13,14^. The BayesS model also allows us to estimate the SNP-based heritability and polygenicity (the proportion of SNPs with nonzero effects) to better describe the genetic architecture for a trait. The application of BayesS has been restricted to GWAS samples with individual-level genotypes but for most common complex diseases, only summary-level data are available. Moreover, despite the implementation of parallel computing strategy^12^, it remains computationally challenging to run BayesS for a biobank-scale data set, as the computing resource increase linearly with the number of individuals or SNPs.

In this study, we enhanced the BayesS model such that the analysis only requires GWAS summary statistics for the trait of interest and a sparse linkage disequilibrium (LD) correlation matrix from a reference sample. Our new method (referred to as Summary-data-based BayesS or SBayesS) opens an unprecedented opportunity to simultaneously estimate the genetic architecture, the parameter *S* (signature of natural selection) and joint SNP effects using publicly available data sets of the largest sample sizes to date. Given the GWAS summary statistics and the sparse LD matrix, it only takes a few hours for analysis with over a million SNPs regardless of the discovery sample size, merely a small fraction of the computational resource required for BayesS. Furthermore, we incorporated functional genomic annotations into the analysis by allowing the distributions of SNP effects to be different among annotation categories (e.g., coding, regulatory, and conserved regions). Compared to other methods utilising functional annotations, such as S-LDSC^15^, BayesRC^16^ and RSS-E^17^, a unique feature of the annotation-stratified SBayesS (referred to as SBayesS-strat) is that it allows the estimation of *S* in each specific functional annotation category. We performed extensive analyses to benchmark between SBayesS and BayesS, and applied the SBayesS methods to GWAS summary statistics from the full release of the UK Biobank^18^ (UKB) data and other published studies^19–27^, followed by time-forward simulations^28^ to facilitate interpretation of the results.

## Results

### Method overview

BayesS is a method that can estimate three key parameters to describe the genetic architecture of complex traits by a Bayesian mixed linear model^12^, namely SNP-based heritability 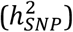, polygenicity (*π*) and the relationship between MAF and effect size (*S*), all of which are defined with respect to a certain set of SNPs (see the definition of 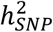 as an example^5^). SBayesS is an extension of BayesS which only requires GWAS summary statistics of the SNPs, and LD information from a reference sample. When the LD values are computed using all SNPs in the GWAS sample, SBayesS model is a linear transformation of the BayesS model without loss of information (Methods), in which case the two models are equivalent in terms of posterior inference (Supplementary Note and Supplementary Fig. 1). However, it is impractical to store pairwise LD correlations of all genome-wide SNPs in the computer memory and not always feasible to access individual-level genotypes of the GWAS sample. We propose to compute pairwise LD correlations between SNPs located on the same chromosome from a reference sample and remove correlations that can be attributed to sampling variation by a chi-squared test, resulting in a sparse LD matrix (Methods). In this case, SBayesS becomes an approximation to BayesS. Assuming the LD reference sample is a random draw from the same population of the GWAS sample, the discrepancy between SBayesS and BayesS arises from the sampling variance of LD correlations used in SBayesS. Ignoring the sampling variance of LD estimates may cause a failure to converge in the Markov chain Monte Carlo (MCMC) sampling process or a bias in parameter estimation (Supplementary Note). In this study, we model analytically the sampling variance of LD estimates as part of the residual variance and allow the estimate of residual variance to vary across SNPs (Methods). Compared to BayesS, SBayesS does not only address the barrier of data sharing as it does not require individual-level data but also substantially increases the computational efficiency because of the use of sparse LD matrix and a different updating strategy in the MCMC sampling (Supplementary Note). These features allow SBayesS to be scalable to data with millions of SNPs regardless of the discovery GWAS sample sizes. In our GCTB software (URLs) where SBayesS is implemented, we have developed a parallel computing strategy to facilitate the computation of the LD matrix. Once the LD matrix is computed, it can be repeatedly used in the analysis of multiple traits. To examine the convergence of MCMC, we use the GCTB-SBayesS implementation of the Gelman-Rubin statistic^29^ which compares the variation between and within multiple chains with different starting values of the model parameters (Methods). Convergence is only concluded if all the three key parameters converge, which may not occur if the LD matrix from a reference sample is too divergent from that of the GWAS sample, or if the summary statistics are generated from a GWAS with low power or contain uncorrected population stratification, poor imputation or other errors such as misreported per-SNP sample size and allele frequency.

To better understand the variability of regional genetic architecture in different parts of the genome, we incorporate functional genomic annotations into SBayesS to allow the three key parameters to vary in different annotation categories (Methods). The functional annotations, such as coding, regulatory and repressed regions, were obtained from the LDSC baseline model^14^ (URLs), where the majority of SNPs (79%) have more than one annotation. To account for the overlap between annotation categories, a SNP that has multiple annotations is assumed to have a mixture of effect distributions, each specific to an annotation category, with the mixing probability *a priori* being one divided by the total number of annotations at the SNP. Note that the effect distribution for an annotation itself is a mixture distribution according to the BayesS prior. Thus, the annotation-stratified SBayesS is a double mixture model. During MCMC sampling, the enrichment of a parameter in an annotation category is computed as the ratio of the sampled value of the parameter in the category to that in the whole genome (Methods). In addition to per-SNP heritability, polygenicity and *S*, we are also interested in the enrichment of per-nonzero-effect (per-NZE) heritability (defined as 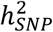 divided by the number of nonzero effects in the category), which is helpful to understand whether the enrichment of 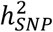 in a genomic region is due to the larger number of associated variants or the larger magnitude of effect size compared to average.

### Benchmarking SBayesS with BayesS

We used the UKB data to evaluate the performance of SBayesS by comparing the results to those from BayesS for common variants. We ran both SBayesS and BayesS with ~1.1 million HapMap3 SNPs with MAF ≥ 0.01 for 18 quantitative traits as analysed in Zeng et al.^12^ (n>100k). We used the HapMap3 SNPs as they were optimised to tag common genetic variants^30^ and are widely used in the literature which improves the comparability of our results with those generated using published GWAS summary statistics. For ease of computation, we used unrelated individuals of European ancestry from the interim release of the UKB data for the BayesS analysis (maximum n=120k across traits) and the same data to generate GWAS summary statistics for the SBayesS analysis. In a random sample of 50k unrelated individuals from the full UKB cohort (n=350k), we computed a sparse LD matrix for the HapMap3 SNPs with a threshold chi-squared value of 10, equivalent to a *r*^2^ threshold of 2×10^-4^ given the sample size (Methods). This resulted in each SNP, on average, being in LD with ~1,000 SNPs on the same chromosome. We show in Fig. 1 that the consistency between the SBayesS and BayesS estimates for all of the three genetic architecture parameters is high across traits (Pearson correlation *r*=0.998 for 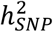, 0.985 for *π* and 0.965 for *S*). The posterior standard error (p.s.e.) of 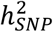 was smaller than that of *π* or *S* in both SBayesS and BayesS because both *π* and *S* are higher-order parameters in the model (Methods).

**Figure 1.**
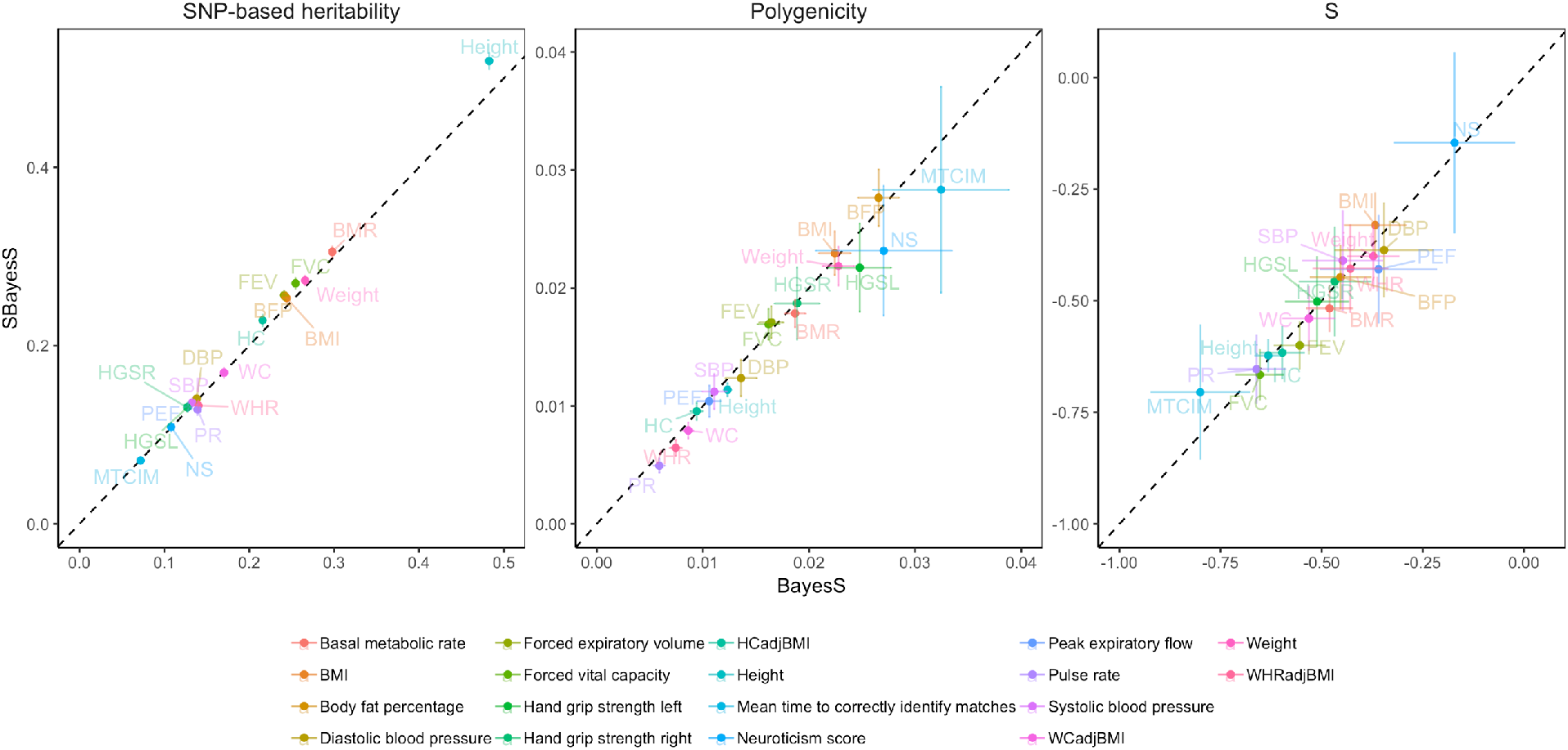
Benchmarking SBayesS with BayesS using the same data for 18 UKB traits. Three genetic architecture parameters were compared, i.e., SNP-based heritability, polygenicity and S, based on the unrelated individuals of European ancestry in the interim release of the UKB data (max n=120k) and ~1.1 million HapMap3 common SNPs (MAF>0.01). The sparse LD matrix used in SBayesS was computed from a random sample of 50k unrelated individuals from the full UKB cohort at a chi-squared threshold of 10 (corresponding to a LD *r*^2^ threshold of 2×10^-4^). Each bar represents the posterior standard error of the estimate. Colours with acronyms indicate different traits, whose full names are shown at the bottom of the figure.

We performed additional sensitivity analyses to investigate the impact of the chi-squared threshold used to filter LD, the SNP panel, the choice of reference sample and the reference sample size on the performance of SBayesS. While a chi-squared threshold of 10 was chosen in this study to balance the true nonzero LD correlations and noise in making the sparse LD matrix (Methods), SBayesS was robust to different chi-squared thresholds used for LD filtering because the SBayesS model accounts for sampling variance of the estimated LD correlation (Supplementary Fig. 2). Compared to the analysis using UKB Axiom array SNPs, the analysis using HapMap3 SNPs tended to give slightly lower estimates of SNP-based heritability and polygenicity but stronger signals of *S* for both SBayesS and BayesS (Supplementary Fig. 3), possibly due to the under-representation of low-frequency SNPs in HapMap3 panel in comparison with the UKB Axiom array panel (Supplementary Fig. 4). Nevertheless, SBayesS was always consistent with BayesS regardless of the SNP panel used (Supplementary Fig. 3). Regarding the LD reference, when the LD reference sample size decreased from 50k to 20k, the differences in parameter estimates were negligible (Supplementary Fig. 5). When the LD reference sample size further decreased to 4k, we observed notable inflation in the estimates of 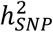 and *S*, suggesting that the LD reference sample size cannot be too small relative to the GWAS sample size (Methods). Given a constant reference sample size (n_ref_=50k), we ran GWAS with sample sizes n_gwas_=120k and 350k and found good concordance between SBayesS and BayesS in both cases (Supplementary Fig. 6). As expected, the *π* estimate from either SBayesS or BayesS increased when a larger ngwas was used because of the increased power to detect small effects. Furthermore, with both n_ref_=50k and n_gwas_=300k held constant, we benchmarked BayesS and SBayesS in a few different scenarios where the LD reference was a subset of the GWAS sample, an independent sample from the same population, or an independent sample from a slightly different population (i.e., the Genetic Epidemiology Research on Adult Health and Aging (GERA) cohort; URLs). When the GWAS and LD reference samples were from the same population, the differences between BayesS and SBayesS were negligible (Supplementary Fig. 7), suggesting that the performance of SBayesS is almost independent of the overlap between the GWAS and LD reference samples as long as they are from the same population. When the GERA cohort was used as the reference sample for the UKB GWAS data, a small inflation was observed in the estimates of S, likely because of the difference in ancestry between the UKB and GERA. This observation demonstrates the importance of choosing a reference sample that is genetically as close to the GWAS sample as possible in the analysis of summary data. Finally, we tested the method in application to ascertained case-control data by simulation. The parameter estimates were nearly unbiased regardless of whether cases were oversampled, although the sampling variances of the estimates of polygenicity and *S* were relatively large in some simulation scenarios where the number of cases was relatively small (Supplementary Fig. 8).

### Analyses of GWAS summary data from the UK Biobank and other published studies

We applied SBayesS to analyse the full release of the UKB data (URLs), including 26 complex traits and 9 common diseases (Supplementary Table 1). Although individual-level data are available in the UKB, application of the standard BayesS to ~350k unrelated individuals with ~1.1 million HapMap3 SNPs would require ~1.5TB memory and ~420 hours with 24 cores. Running SBayesS only requires approximately 15GB memory and 8 hours with 4 cores, demonstrating the resource-efficiency and scalability of SBayesS. Prior to running SBayesS, we carried out standard quality control (QC) of the data (Methods) and used linear regression to perform a GWAS analysis in unrelated individuals to generate summary statistics for each trait. We also applied SBayesS to data for 9 other complex common diseases from published GWAS of very large sample size where only summary statistics are available (Supplementary Table 2). In the analysis of the UKB data, we used the sparse LD matrix computed from 50k individuals (a random subsample of all the UKB unrelated individuals as described above). For the analysis of data from published GWAS of which nearly all the samples are of European ancestry, the GERA sample was used as the LD reference. To mitigate the problem due to inconsistent LD between the GWAS and reference samples, we excluded SNPs in the major histocompatibility complex (MHC) region. The SNP-based heritability estimates for the diseases were converted to those on the liability scale following the method in Lee et al^31^.

On average across the 44 complex traits (including diseases), 1.8% of the 1.1 million common HapMap3 SNPs explained 18% of the phenotypic variance (Fig. 2 and Supplementary Table 1-2). The estimate of SNP-based heritability for height was 0.545 (p.s.e.=0.003), consistent with those in previous studies using different approaches and data sets^6,13,32–34^. The most polygenic traits (i.e., body fat percentage, educational attainment and schizophrenia) had about 5% (~55,000) SNPs with nonzero effects. The least polygenic traits were prostate cancer, age at menopause and male pattern baldness, which were affected by about 0.1-0.3% (1,000-3,000) common SNPs. The estimate of *S* was significantly negative (*P*<0.001) in all the traits analysed, suggesting a pervasive action of negative selection on the trait-associated variants. Interestingly, most traits had an estimate of *S* at about −0.6 (median=-0.578, SD=0.096), which motivated us to further investigate the interpretation of *Ŝ* with respect to natural selection (see below). There were some traits for which *Ŝ* was not significant in previous analysis based on individual-level genotypes and array SNPs^12^ (e.g., T2D, fluid intelligence and neuroticism score) but became significant in the current study, likely because of the increased sample size that improved the detection power (Supplementary Fig. 6). We also re-ran the analysis for the 9 public GWAS data sets with the UKB subsample as the LD reference and found that the results were highly consistent with those using LD from the GERA (Supplementary Fig. 9).

**Figure 2.**
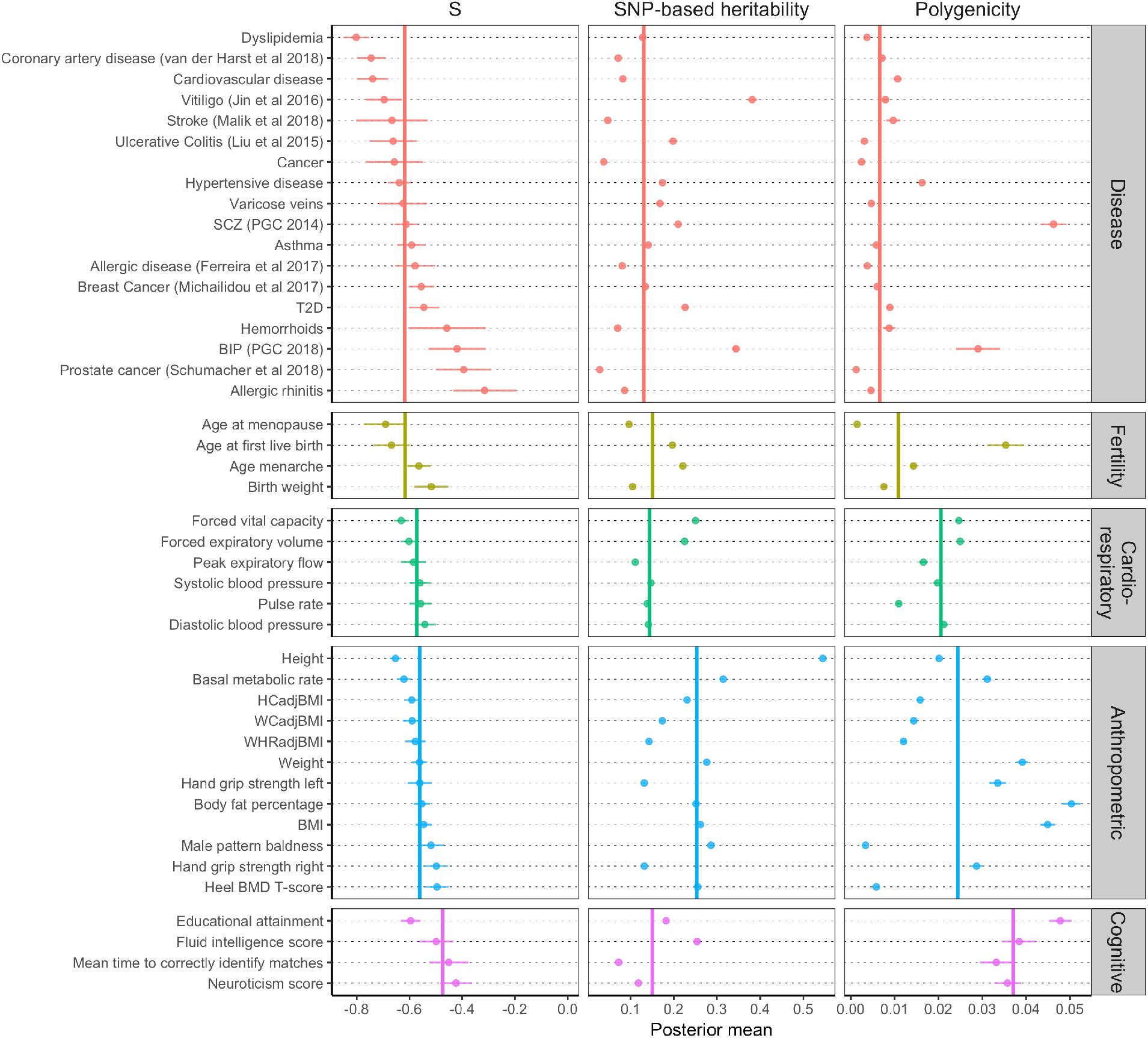
Estimation of the three genetic architecture parameters for 44 complex traits (including diseases) in UKB (max n=350k) and 9 common diseases from published GWAS (labelled with publications). Shown are the posterior means (dots) and standard errors (horizontal bars) of the parameters for each trait. The colour indicates the category that the trait belongs to. The vertical bar shows the median of the estimates across traits in each category.

We broadly classified the 44 traits into five categories related to disease, fertility, cardiorespiratory, anthropometry and cognition. The estimates of the genetic architecture parameters varied across traits and appeared to have distinct patterns in different categories (Fig. 2). Anthropometric traits had a substantially higher median SNP-based heritability (0.253) than the other categories, among which the differences were small (0.097–0.151). The median value of the polygenicity estimates was the lowest for diseases (0.007) and the highest for cognitive traits (0.037). The estimates of polygenicity for psychiatric disorders such as schizophrenia (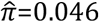, p.s.e.=0.003) and bipolar disorder (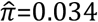, p.s.e.=0.009) were substantially higher than that for other types of disease and comparable to those for the cognitive traits, consistent with the high polygenicity for brain-related traits reported in previous studies^10,35^. The estimate of |*S*| was the highest for diseases, especially cardiovascular disease, and fertility traits, and the lowest for cognitive traits, with a relatively large variability in *Ŝ* for diseases.

To investigate the diversity of genetic architecture in more traits, we applied SBayesS to GWAS summary data from the Neale Lab (URLs) for 274 quantitative traits in the UKB, among which 130 passed the convergence test (the failed ones were mainly due to the smaller sample sizes; mean n = 231k for converged and 73k for not converged) and 110 were not included in the analyses above (Supplementary Table 3). Fig. 3 shows the distributions of the estimated genetic architecture parameters for the total 155 traits including 137 complex traits and 18 common diseases in the UKB and published GWAS. Similar to that for the 44 traits analysed above, the distribution across traits was relatively flat for 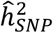, skewed to the right for 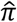, and symmetrically distributed with a large mass around −0.6 for 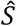. Among the 155 traits, 79% had 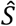 within the range of −0.7 to −0.5 (median=-0.576). In addition, *Ŝ* was weakly correlated with 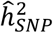 (Pearson’s correlation=-0.191), and the variation of *Ŝ* decreased with increasing 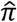, suggesting some interrelationship among the genetic architecture parameters. Next, we investigated the interplay of genetic architecture parameters under natural selection through simulation.

**Figure 3.**
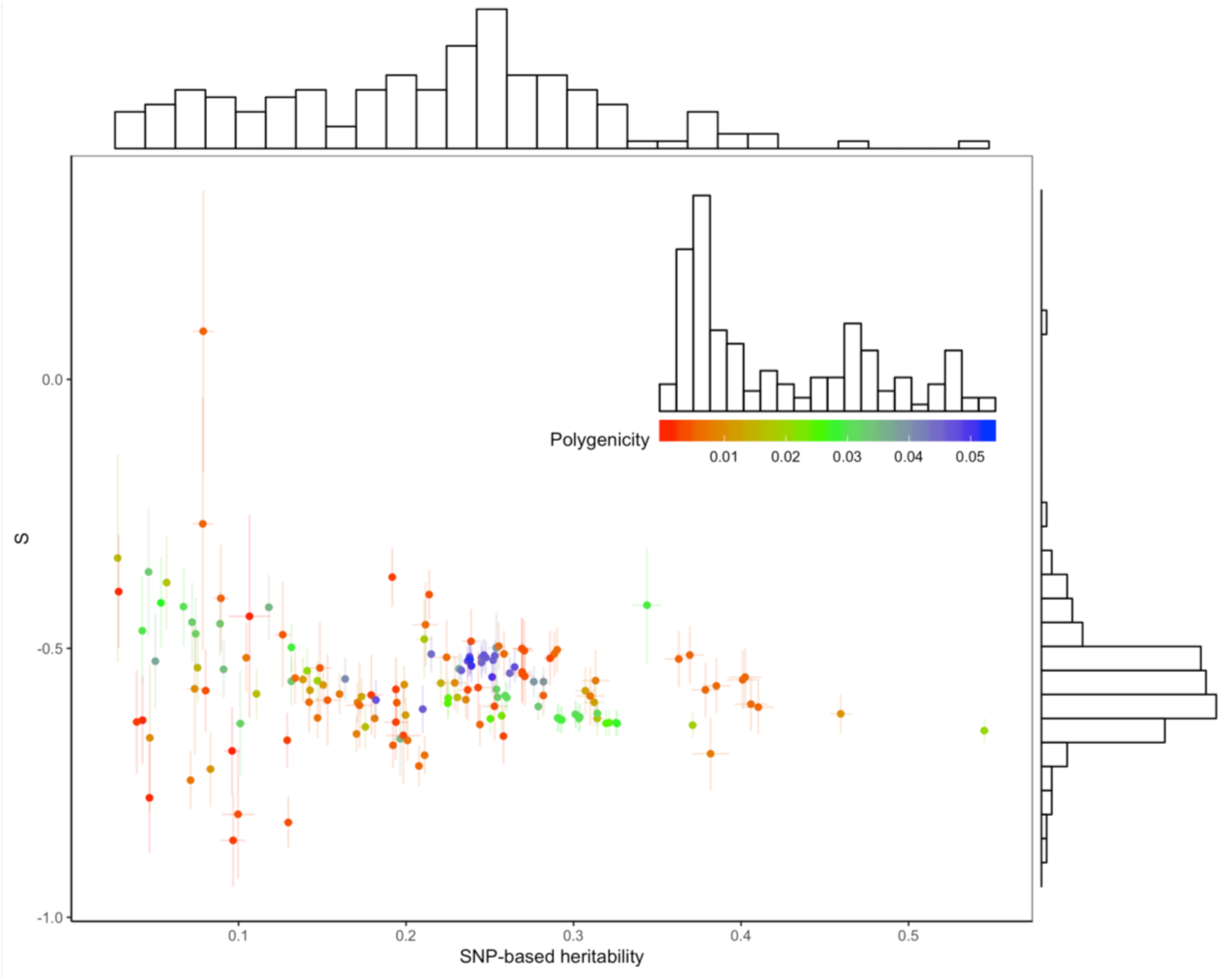
Estimation of the genetic architecture parameters for 155 complex traits (summary data for 130 quantitative traits from the Neale Lab and 25 traits and diseases from our GWAS analyses and published studies). The estimated S is plotted against the estimated SNP-based heritability with the corresponding posterior standard errors (bars) as well as the marginal distributions of the estimates. Colour indicates the estimate of polygenicity for each trait where the scale and distribution are shown in the inset graph.

### Projecting the real data analysis results onto the genetic architecture patterns observed from simulations

Although a negative estimate of *S* is a signature of negative selection, the numeric interpretation of *Ŝ* is still not clear. For example, our results showed that most traits had *Ŝ* at about −0.6; does it mean that negative selection acted on the associated variants with similar selection strength among traits? To answer this question, we performed forward simulations^28^ given the mutational target size of the genome (*π_m_*; the proportion of DNA sequence that can produce mutations affecting the trait) and total trait heritability at all the mutations (*h*^2^). Note that *π_m_* and *h*^2^ are the evolutionary parameters underlying *π* and 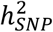. The simulations were based on a demographic model inferred by Gravel et al^36^. A normalising stabilising selection model^37^ was used to link phenotype to individual fitness with a given selection strength (Methods). In the last generation of selection, we computed the genetic architecture parameters 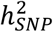, *π* and *S* at all the common causal variants (Methods). Repeating the simulation with different values of *π_m_*, *h*^2^ and selection strength produced a landscape of the genetic architecture under different scenarios (Fig. 4). When there was no selection (selection strength = 0), 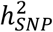 and *π* closely reflected the values of *h*^2^ and *π_m_*, respectively, with *S*=0. For given values of *h*^2^ and *π_m_*, increasing selection strength shaped the genetic architecture on three sides. First, *π* became smaller than *π_m_* because a larger fraction of trait mutations was kept at MAF<0.01 due to the action of negative selection. Second, a gap between 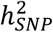 and *h*^2^ was present as a result of excessive rare trait mutations (MAF < 1%), one of the explanations to the missing heritability problem^38^. Third, the magnitude of *S* increased, indicating a stronger negative relationship between effect size and MAF than that under weaker selection given constant *h*^2^ and *π_m_*. It is important to note that a high |*S*| could be achieved even with a relatively low strength of selection, when per-variant heritability (in proportion of *h*^2^/*π_m_*) was high, suggesting that we cannot infer the strength of natural selection based only on *S* without taking the other components of the genetic architecture into account. Moreover, the same level of polygenicity (*π*) could be observed across different values of mutational target size and strength of negative selection, indicating that polygenicity is driven by both factors rather than negative selection alone (note that O’Connor et al^10^ concluded that negative selection is responsible for polygenicity, which was defined depending on the number of large effects).

**Figure 4.**
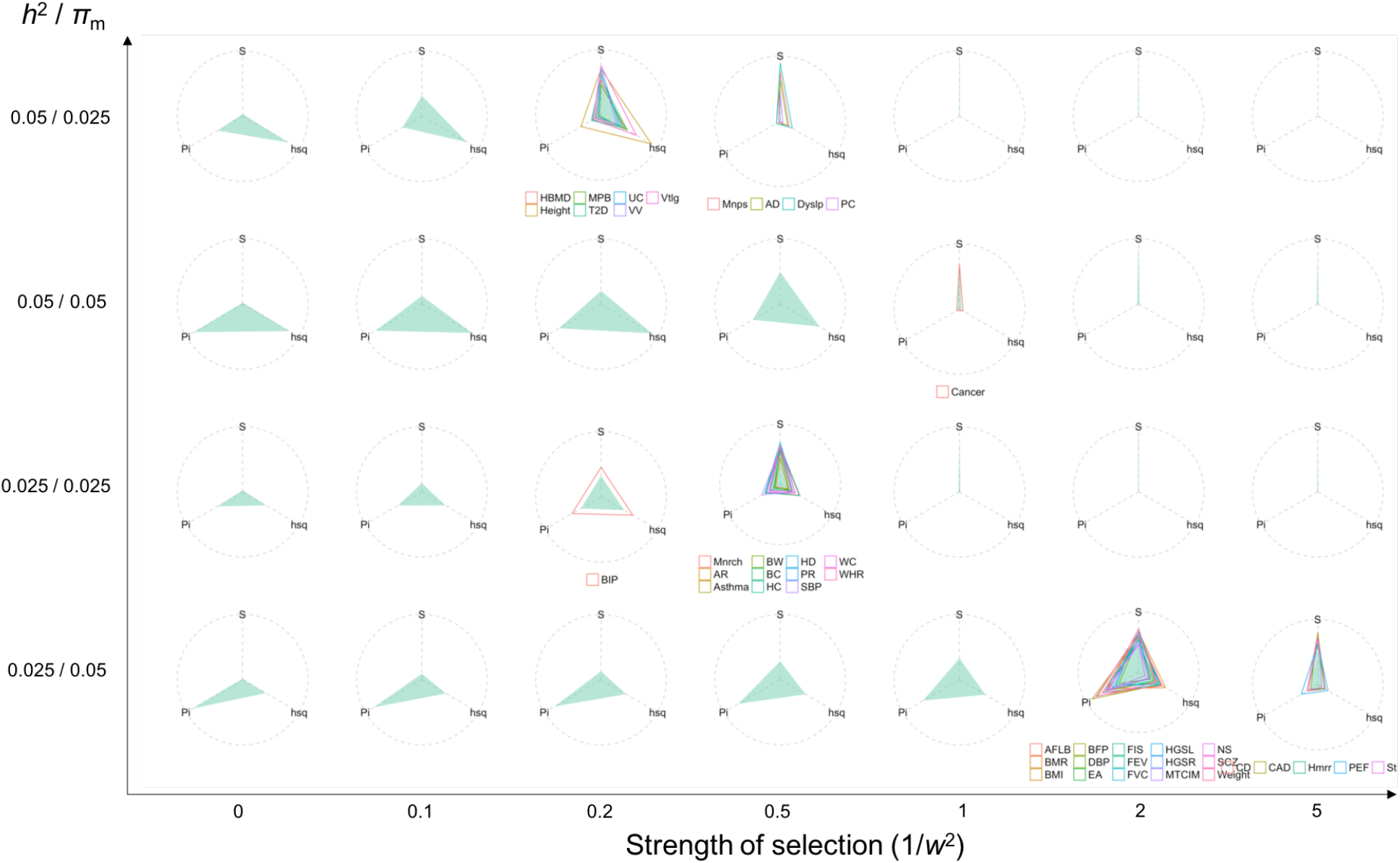
Projection of real data analysis results onto a map of genetic architecture patterns observed in simulations. Both x and y axes are input values in the simulation, where x-axis is the strength of selection denoted as one over the variance (*w*^2^) of the phenotypes surrounding at the optimum fitness value (a classic model for stabilizing selection), and y-axis is the ratio of the heritability at all causal variants (*h*^2^) over the proportion of mutational targets (*π_m_*), which is proportional to the per-mutation heritability. Each circle consists of the values of SNP-based heritability, polygenicity and *S*, scaled by the maximum values across simulated or real traits such that the three parameters are on the same scale from zero to one. The green shadow indicates the computed values of the three genetic architecture parameters at the common causal variants (MAF>0.01) in the last generation of the forward simulation given the input values on x and y axes. The hollow triangle shows the estimated genetic architecture for each UKB trait. A demographic model proposed by Gravel et al (2011) was used in the simulation.

Given the evolutionary parameters used in the forward simulation, we observed a variety of patterns of genetic architecture, which can be used as a map to infer the underlying evolutionary parameters in real traits. We then projected the observed genetic architecture of the 44 complex traits and diseases onto the map generated from the forward simulation (Fig. 4). The projection was done by minimizing the sum of the differences in internal angles between the two “triangles” (the observed versus the simulated genetic architecture) (Methods). Our projection shows that although the variation of *Ŝ* was small among traits, different traits had diverse strength of negative selection on the associated variants. No trait was projected to the bottom left of the map where both per-variant heritability and strength of selection were relatively low, or the top right of the map where the selection was so strong that most causal variants were rare or had been removed from the population. As a result, there was a negative correlation between per-variant heritability and strength of selection from the projected genetic architecture. This inverse relationship is reasonable given *S*, because variants of larger per-variant heritability are expected to receive higher selection pressure such that a strong signature would manifest itself even if the selection on the trait is not as strong as that on a trait with a higher polygenicity and/or lower heritability. According to the projection, SNPs associated with brain-related traits including cognition and psychiatric disorders were under relatively strong selection, and most common diseases tended to have relatively small mutational target size. When we projected the median estimates of the genetic architecture parameters for each trait category onto the map, we found relatively strong selection on genetic variants associated with cardiorespiratory and cognitive traits and relatively small mutational target size for diseases and fertility traits (Fig. 5a). As a result, common variants associated with diseases showed the largest estimated effect variance, and those associated with cardiorespiratory and cognitive traits showed the smallest estimated effect variance (Fig. 5b), consistent with the GWAS results where the average marginal effect size (in SD units) decreased from disease-to cognition-associated SNPs (Fig. 5c,d). We further performed projections based on the simulations with a constant effective population size of 10,000 and did not observe a qualitative change on the conclusions (Supplementary Fig. 10 and 11), suggesting a limited impact of demography on the formation of genetic architecture.

**Figure 5.**
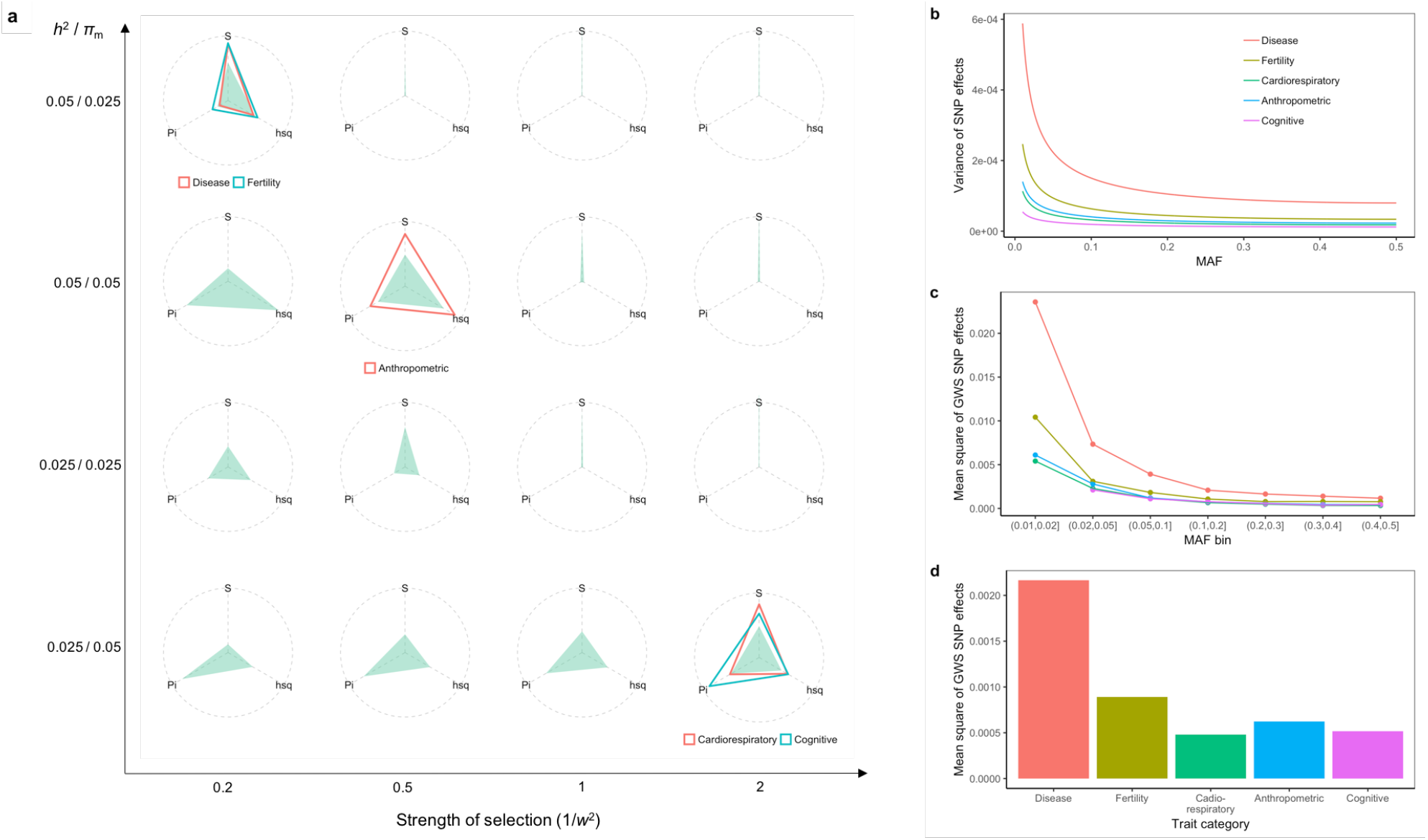
Differentiated signatures of negative selection in the genetic architecture of different trait categories. a) Projection of observed genetic architecture averaged over each trait category onto a map of patterns from forward simulation. b) The expected variance of SNP effect as a function of MAF based on the model 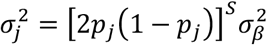, with *Ŝ* and 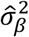. being the mean estimates from SBayesS across traits for each category. c) The mean squared effect size as a function of MAF for genome-wide significant (GWS) SNPs from GWAS in the 5 trait categories. d) The overall mean squared effect size of the GWS SNPs regardless of MAF. In b-d), The SNP effects are in standard deviation units of liability for diseases or phenotype for quantitative traits.

### Analyses incorporating functional genomic annotations

The functional annotation categories used in our analysis were from the LDSC baseline model^14^. We excluded continuous annotations and annotations with flanking windows, resulting in 21 annotation categories such as the coding, regulatory, repressed and conserved regions (Supplementary Table 4), with a large proportion of overlap between categories (Supplementary Fig. 12). We applied SBayesS-strat to the 35 UKB traits (including 9 diseases), and combined the parameter estimates across traits for each functional category based on a method that accounts for the phenotypic correlations between traits (Methods). In general, the fraction of 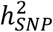 explained by each functional category was proportional to the size of the category (correlation=0.987; Supplementary Fig. 13). Repressed regions appeared to be an outlier with the estimated 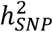 significantly lower than its expectation. The correlation was even higher between the polygenicity and the size of functional category (correlation=-0.999), suggesting that a functional category that explains a greater fraction of heritability tends to have a larger number of causal variants, consistent with the findings of O’Connor et al^10^ with a different approach. The average value of *Ŝ* was −0.607, ranging from −0.662 in repressed regions (s.e.m.=0.010) to −0.555 in transcription start sites or TSS (s.e.m.=0.032).

To better distinguish the contributions of the number and the magnitude of the nonzero effects to 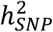, we estimated per-NZE heritability 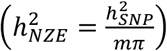 in each category, and computed the fold enrichment by comparing the per-category estimate to the genome-wide estimate (Methods and Supplementary Table 5). The per-NZE heritability showed the highest enrichment at TSS, followed by coding, 5’-UTR, 3’-UTR and conserved regions, and remarkable depletion in the repressed regions (Fig. 6a). In addition, the enrichment in per-NZE heritability was proportional to the enrichment in polygenicity (correlation=0.982), suggesting that the larger per-SNP heritability in a functional category was not only because of the larger number of causal variants but also the larger effect sizes, confirmed by forward simulation (Supplementary Fig. 14). The parameter *S* was enriched in the repressed, coding, 5’-UTR and conserved regions despite a large sampling variation in the estimated fold enrichment (Fig. 6b). There was a negative correlation of −0.290 between the enrichments of *S* and per-NZE heritability, and the correlation decreased to −0.788 when the apparent outliers (i.e., the coding, 5’-UTR and conserved regions) were removed. The presence of a negative correlation could be because for biologically important regions, a fraction of the genetic variants has been under positive selection, which to some extent offsets the signature of negative selection. To test if the negative correlation was driven by any artificial effect, we applied the method to simulated data sets under the null model (no enrichment in genetic architecture parameters) and did not observe a significant correlation (Supplementary Fig. 15). Although the coding and repressed regions had similar *Ŝ*, the variance of effect sizes were different regardless of MAF (Fig. 6c,d), suggesting different distributions of effect sizes for SNPs in different functional genomic regions.

**Figure 6.**
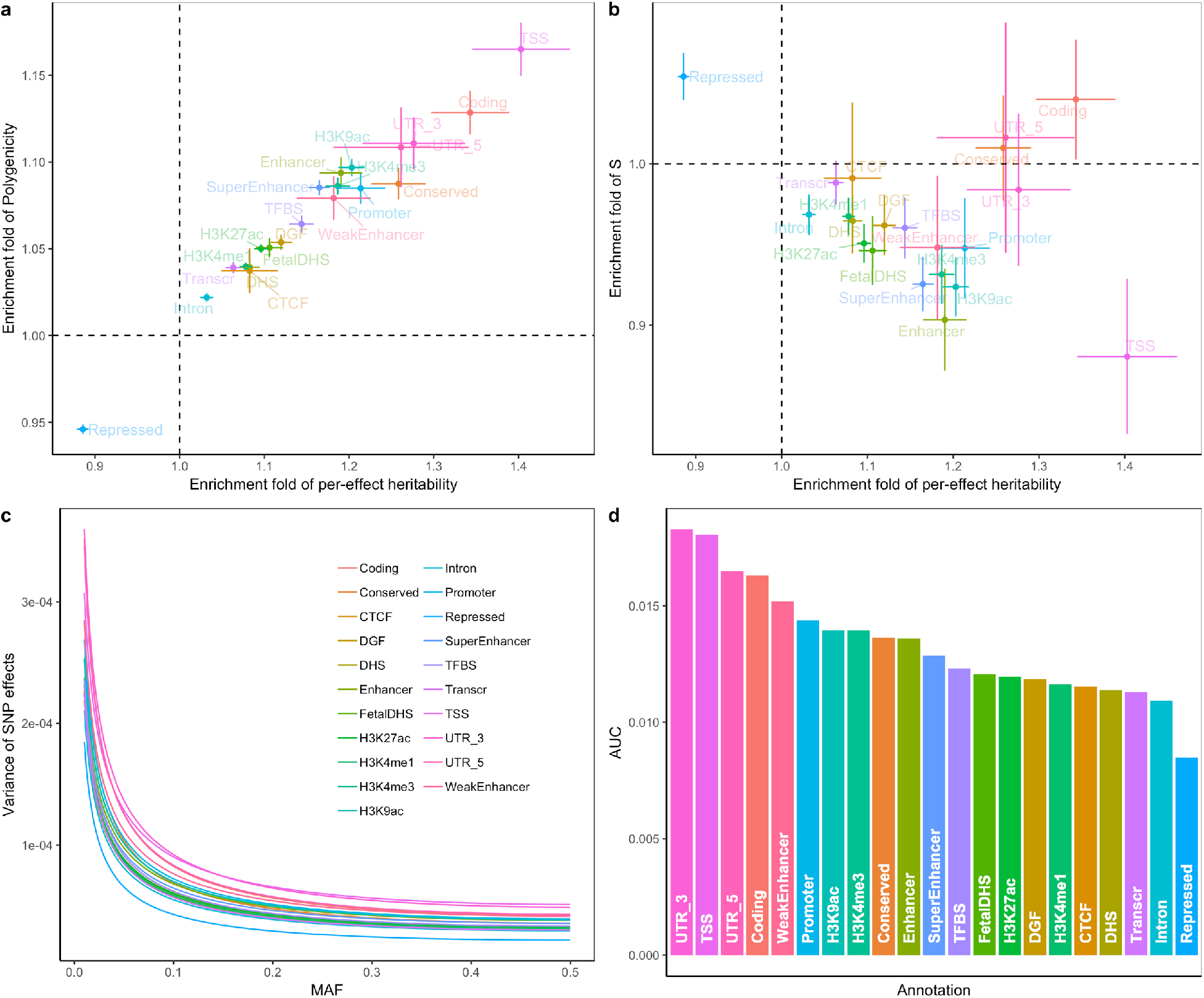
Characterisation of the genetic architecture in 21 functional genomic annotation categories using the SBayesS-strat model. a) Fold enrichment of per-NZE heritability against that of polygenicity. b) Fold enrichment of per-NZE heritability against that of *S*. Each dot is the mean across 35 UKB traits (including diseases). Each bar indicates the estimated standard error of the mean. c) Shown is the expected variance of effect sizes as a function of MAF based on the model 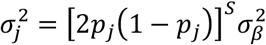, with *Ŝ* and 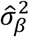. being the mean estimates across traits for each annotation category. d) The area under the curve (AUC) of the expected effect variance over MAF for each annotation category, as shown in c).

## Discussion

We have developed an efficient summary-data-based method to estimate the joint distribution of effect sizes and MAF as well as SNP-based heritability, polygenicity and joint SNP effects. By analysing GWAS summary statistics from the public domain, we detected pervasive signatures of negative selection in the genetic architecture of a wide range of complex traits including common diseases (Fig. 2 and 3). This means, assuming that most mutations are deleterious to fitness, mutations with larger effects on fitness are more likely to be eliminated or kept at lower frequencies in the population by negative selection. Interestingly, most traits had *Ŝ* at about −0.6 with diverse estimates of 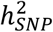 and polygenicity, implying that the *S* = −1 model originally used in the GREML method^39^ is more appropriate than the *S* = 0 model for most complex traits. Schoech et al^7^ linked the *S* parameter (denoted as *α* in their infinitesimal model using profile maximum likelihood estimation with individual-level data) to the *τ* parameter in Eyre-Walker’s model^3^ and further drew inference on the average genome-wide selection coefficient. However, our forward simulations have shown that inference regarding the strength of selection cannot solely be made based on *S* but should take other genetic architecture parameters into account. Our projection results show that despite the narrowly distributed *Ŝ*, the strength of selection varied substantially across traits (Fig. 4), assuming stabilising selection in action (as suggested by previous work^11,40,41^). In general, diseases showed the smallest mutational target sizes, while cardiorespiratory and cognitive traits showed the strongest selection on the associated variants (Fig. 5a). As a result, common variants associated with diseases (heart, lung, and brain function) presented the largest (smallest) effect sizes (Fig. 5b-d).

Our annotation-stratified analysis further revealed that negative selection has acted differentially on the genome, resulting in non-identical distributions of effect sizes between functional categories. The biologically important categories, such as the TSS, coding and 5’-UTR regions, had the highest enrichment in per-NZE heritability and polygenicity, whereas the repressed regions were depleted in both per-NZE heritability and polygenicity (Fig. 6). Previous studies have attributed the enrichment in per-SNP heritability to the larger number of causal variants in functional regions^10^. Here, we further highlighted the contribution of differences in effect sizes to the per-SNP heritability enrichment. The repressed regions were evidently enriched in *Ŝ* and had smallest effect variance for common SNPs (Fig. 6), suggesting an apparent role of negative selection in constraining the frequency of mutations with large effect sizes in the repressed regions. This observation should not be too surprising given that chromatin repression represents a higher order functional domain and is enriched for evolutionarily conserved non-exonic sequences^42^. Coding and 5’-UTR regions were enriched for *Ŝ* yet had largest effect variance for common SNPs. Our explanation is that in addition to negative selection, positive selection may have also played a role in these regions to drive the evolution of protein-coding genes such that SNPs with relatively large (favourable) effects could be common. Unfortunately, our method cannot distinguish negative from positive selection when both types of selection are in action.

There are several limitations in this study. First, we used HapMap3 common SNPs for analysis and extrapolated our inference on negative selection to unobserved rare variants. We expect that the signature that natural selection left in common variants would remain in rare variants. Because the rare sequence variants are not fully available in the UKB, we investigated this hypothesis by forward simulation. As expected, the selection signature in rare variants followed similar patterns as that in common variants, but the magnitude of *S* was weaker (Supplementary Fig. 16). This is because the very rare variants were mostly new mutations whose relationship between effect size and MAF had not yet been shaped by selection, which diluted the selection signals from the variants under selection (Supplementary Fig. 17), suggesting that the true *S* parameter is allelic age dependent and subject to the combined effect of mutation, selection and genetic drift. An apparent change in the effect size-MAF relationship when moving toward low MAF was also reported by Schoech et al^7^. Second, we projected the parameter estimates onto the patterns observed from evolutionary simulations under a model of stabilising selection with a constant trait heritability. A violation of the assumption that only stabilising selection is in action could complicate the comparison of evolutionary parameters between traits. For example, according to our projection, genetic variants associated with BMI are under stronger negative selection than those associated with height, which may not be true if there is a relatively strong positive selection in height (see Supplementary Fig. 18 for a simulation study). The strong negative selection on BMI-associated variants inferred from our result is nevertheless consistent with a recent study^43^ which showed a large proportion of variance explained by very rare variants (0.01%<MAF<0.1%) for BMI although the SE was large. We also performed a set of simulations with fixed environmental variance allowing the heritability to reach a mutation-selection-drift equilibrium. The results (Supplementary Fig. 19 and 20) were comparable to those from the simulations with fixed heritability and variable environmental variance presented above (Fig. 4 and 5a). Third, independence of chromosomes is assumed in our model. This may not hold if there was non-random mating in the ancestral population. For example, assortative mating would introduce positive correlations between trait-increasing alleles located on different chromosomes, and therefore increase heritability in the equilibrium population, e.g. for height^44^. Our estimate of SNP-based heritability accounts for intra-chromosomal LD but ignored inter-chromosomal LD so that it is expected to be between the base and equilibrium population estimates. Further improvement is possible if the correct correlation matrix is used. Despite these limitations, our study highlights the impact of negative selection on the genetic architecture across complex traits and in different functional genomic regions. In addition to better understanding of the genetic architecture, our methods can also be applied to genetic mapping and polygenic risk prediction through the use of the joint SNP effect estimates or the characterised underlying distributions of effect sizes as prior knowledge for other methods^45^.

## Methods

### SBayesS

Let us consider an individual-level data-based multiple regression model for a GWAS cohort:

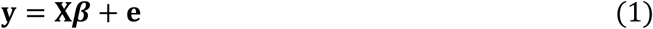

where **y** is the vector of phenotypes adjusted for all fixed effects, **X** is the column-centred genotype matrix, ***β*** is the vector of SNP effects, and **e** is the vector of residuals with 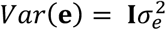 for a cohort of unrelated individuals. Assuming Hardy-Weinberg equilibrium (HWE), the variance of genotype dosage (0, 1, 2) of SNP *j* is *h_j_* = 2*p_j_q_j_*, where *p_j_* is MAF and *q_j_* = 1 − *p_j_*. Let **D** be a diagonal matrix with 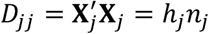, where *n_j_* is per-SNP sample size. Multiplying both sides of the model by **D**^−1^**X**′ gives

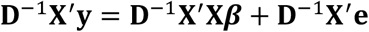

Note that **D**^-1^**X′y** = **b**, the vector of least squares estimates of SNP marginal effects from GWAS, and 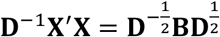, where 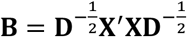 is the LD correlation matrix among all SNPs (ref^46^). Let 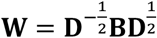 and ***ε*** = **D**^−1^**X′e**. Then, the above equation can be written as

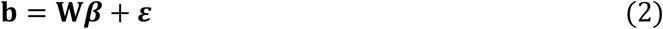

In contrast to the identity structure of residual variance in model (1), the residuals in model (2) are dependent of LD, as

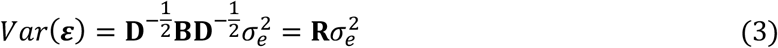

This is a generic form of summary-data-based Bayesian regressions, which is similar to Zhu and Stephens’s RSS model^33^. As in BayesS, we assume the effect size is related to MAF through a parameter *S*:

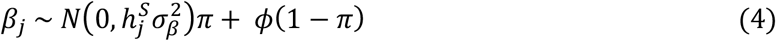

where *ϕ* is a point mass at zero, and *S* (the relationship between MAF and effect sizes), 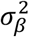 (the effect variance factor common to all SNPs) and *π* (the proportion of SNPs with nonzero effects, i.e., the polygenicity) are considered as unknown, with prior distributions of a standard normal, a scaled inverse chi-squared distribution (Supplementary Note), and a uniform distribution between zero and one, respectively. Specifying a different prior distribution to *β_j_* gives a form of other summary-data-based Bayesian alphabet models^47^.

We show in the Supplementary Note that models (1) and (2) are equivalent in terms of posterior inference. This is because the GWAS estimates of SNP effects (**b**) and LD correlation matrix (**B**) are sufficient statistics for the joint posterior distribution of ***β***, i.e., under BayesS prior (assuming *π*=0 for simplicity),

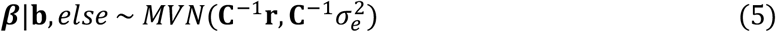

where 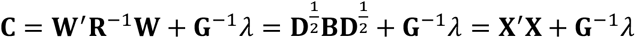 and **G** is a diagonal matrix with 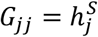 and 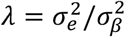, and **r** = **W′R**^−1^**b** = **Db** = **X′y**. Compared to model (1), model (2) allows us to incorporate LD information from a different reference sample from the GWAS sample for which the individual-level data are often not accessible. Further, it is often not practical to compute and store the entire LD matrix in the memory. Therefore, we used a sparse LD matrix that ignores the small LD correlation estimates that are due to sampling variation, but still accounted for the sampling variance of LD correlation in the model (see below). Once the LD matrix is computed, it can be used repeatedly in the GWAS summary-data analysis for different traits.

We used MCMC algorithm to generate 50,000 posterior samples (the first 20,000 discarded as burn-in) from the joint posterior distribution of model parameters, based on which statistical inference was made. Details of the MCMC sampling scheme are shown in the Supplementary Note. The posterior mean was used as the point estimator, with the statistical uncertainty quantified by the posterior variance or its square root (posterior standard error), as shown below. We ran 4 parallel chains with different starting values of the parameters randomly sampled from their prior distributions. Following the method proposed by Gelman and Rubin^29^, we estimated the posterior variance by

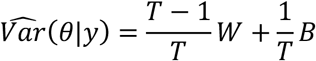

where *T* is the chain length, *W* is the within-chain variance, and *B* is the between-chain variance. To assess convergence in MCMC, we computed the potential scale reduction statistic

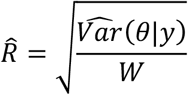

for each of the model parameters. As suggested by Gelman and Rubin, 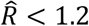 generally indicates good convergence. Thus, we concluded convergence for a trait analysis when all of the three genetic architecture parameters had 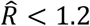.

### Sparse LD matrix

For computational efficiency, we used a sparse LD matrix in the analysis where LD due to sampling variation were set to be zero. To this end, we tested whether the LD between each pair of SNPs on the same chromosome is zero in the population when computing the LD correlation matrix using a reference sample. Under the null hypothesis that the true LD in the population is zero, we assume (ref^48^)

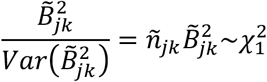

(tilde denotes quantities computed from the reference sample) and reject the null if the chi-squared statistic > 10 (*P*<0.0016). This is equivalent to a *r*^2^ threshold of 2×10^-4^ given a sample size of 50,000. We set 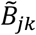 to be zero if the null hypothesis is not rejected or if the two SNPs are on different chromosomes, leading to a sparse LD matrix. The chi-squared threshold of 10 is chosen in order to balance the type I and II error rates. If a type I error occurs, i.e., the true correlation ρ_jk_=0 but 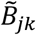 is not set to be zero, then as explained below, 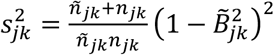, which is very likely to be larger than the true sampling variance of 1/*n*_jk_. This would inflate the residual variance and therefore deflate the heritability estimate. If a type II error occurs, i.e. ρ_jk_≠0 but 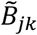 is set to be zero, then 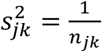, which is very likely to be larger than the true sampling variance of 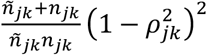. This would deflate the residual variance and therefore inflate the heritability estimate. Since the consequence of type II errors is worse, we use a not-too-stringent threshold to eliminate the LD due to sampling. This also suggests that LD reference sample size cannot be too small, otherwise, type II error rate would increase due to the loss of power. Since we only include non-zero elements in the LD matrix, it is faster by folds to run the summary-level data analysis with substantially less amount of memory needed.

### Modelling LD sampling variance

The use of a sparse LD matrix from a reference sample will result in two sources of sampling variation. The first is the difference in sampling variance between the reference and GWAS samples for the LD correlations included in the sparse LD matrix. The second is the sampling variance of LD correlations that are set to be zero. As shown in the Supplementary Note, ignoring these sampling variations will result in a bias in the mean of the full conditional distribution of *β*_j_ and thereby biases in the estimation of other model parameters. Here, we account for both sampling variations in the model, as described below.

Suppose that the observed LD correlation between SNP *j* and *k* equals to the true population LD plus a deviation, i.e., *B_jk_* = *ρ_jk_* + *δ_jk_*, or in the reference sample, 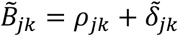. Then, the LD correlation in the GWAS sample is

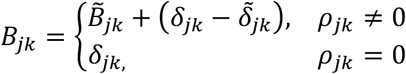

Let Δ_jk_ denote the unobserved quantity above, i.e.,

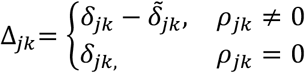

In Eq. (2), we can write

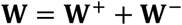

where 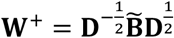 is the observed data, and 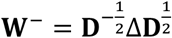 is not observed. Substituting **W** in Eq. (2) by **W**^+^ gives

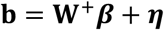

where ***η*** = **W^−^*β* + *ε*** are the new residuals that contain the differences in sampling deviations of LD between GWAS and reference samples when the population LD are not zero, and the sampling deviations of LD in the GWAS sample when the population LD are zero. Conditional on **Δ**, the residual variance is

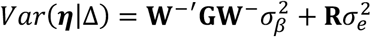

with 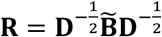. However, this cannot be computed because **Δ** is not observed. As shown in the Supplementary Note, unconditional on **Δ**, the marginal residual variance and covariance are

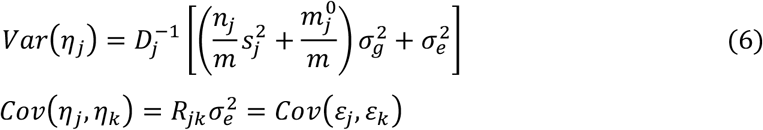

where 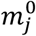 is the number of SNPs not in LD with SNP 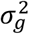 is the trait genetic variance, and

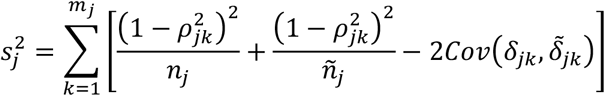

is the total sampling variance for non-zero LD (with *ρ*_jk_ approximated by 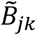 in practice). In the absence of sample overlap between the LD reference and GWAS samples, 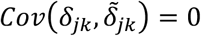. In the case of complete sample overlap, 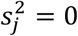. We therefore have the following observations:

1. The LD sampling variance only affects the variance but not covariance of the model residuals. Thus, accounting for the LD sampling variance in the Gibbs sampling of *β_j_* is straightforward, as shown in the Supplementary Note.
2. The LD sampling variation has two components, one due to the use of a different reference sample for LD information and the other due to the use of a sparse LD matrix, both of which are proportional to the genetic variance. If LD are estimated from the GWAS sample, 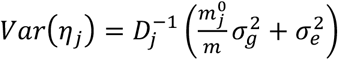. Further, if the genome-wide full LD matrix is used, 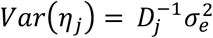, the same as that in Eq. (3).
3. If SNP *j* is independent of all other SNPs, 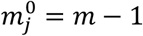 and 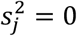. Therefore, 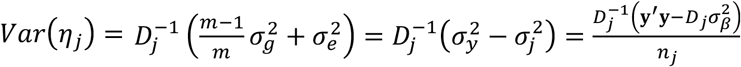 which is the residual variance under a single-SNP GWAS model.
4. Under some conditions, e.g., small *ñ_j_* but large 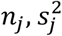 can be greater than 1. Thus, in the presence of LD sampling variance, the total residual variance (in the square brackets of Eq. 6) can be greater than the phenotypic variance of the trait.

### SNP-based heritability estimation

In BayesS^12^, we computed the genetic variance 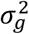 as the variance of genetic values across individuals given the sampled values of ***β*** in each MCMC iteration. As described in Zhu and Stephens^33^, this is equivalent to the following quadratic term of ***β*** given the LD correlation matrix:

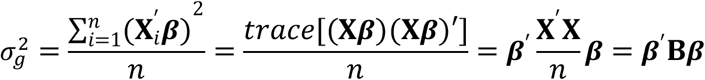

Given the right-hand-side updating strategy in MCMC (Supplementary Note), this quadratic term can be computed efficiently as the difference of two vector-by-vector products:

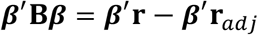

where **r** is defined as in Eq. (5) and **r**_*adj*_ is the adjusted **r** for ***β***. The residual variance 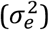 is sampled from a scaled inverse chi-squared distribution with the mean mainly driven by

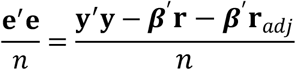

where **y′y** is estimated by the median value of 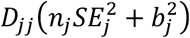 across SNPs, where *SE_j_* is the standard error of *b_j_*. Conditional on 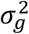 and 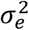, we computed 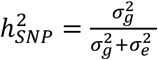 in each MCMC iteration, and used the mean over MCMC samples as the point estimator of the SNP-based heritability.

### Annotation-stratified SBayesS

To accommodate annotation overlaps, each SNP effect is assumed to have a mixture distribution with respect to the functional categories annotated at the SNP. Suppose there are Γ categories in the genome in total, and *K* categories overlapped at SNP *j*. Then, the prior distribution for the effect is

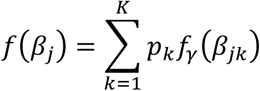

where *p_k_* = 1/*K* is the prior probability with the effect following a distribution specific to the *k*^th^ category (e.g., category *γ* = 1, …, Γ). According to the BayesS prior (Eq. 4), *f_γ_*(*β_jk_*) itself is a mixture distribution

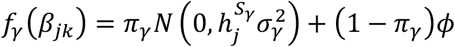

with independent standard normal, scaled inverse chi-squared and uniform priors for 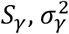 and *π_γ_*, respectively. Our main interest was to estimate the genetic architecture parameters *S_γ_, π_γ_* and 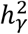 for each category, whose full conditional distributions only depend on the effects of SNPs in the corresponding category. Apart from the per-SNP heritability, we also defined per-nonzero-effect (per-NZE) heritability 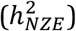 as the total heritability explained in a category divided by the number of nonzero effects in the category. In addition to the category-specific parameters, we estimated the global parameters 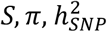 and 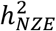. empirically conditional on the sampled value of ***β*** in each iteration of MCMC. The fold of enrichment for each parameter in each trait was then computed as 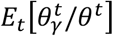 over *T* MCMC iterations. The estimation variation of the enrichment fold was quantified by the posterior variance as described above.

We further combined the information across traits by calculating the mean fold enrichment for each functional category. To account for the phenotypic correlation among the traits, we estimated the effective number of traits (*n_e_*) by performing an Eigen decomposition on the phenotypic correlation matrix, following the method described in ref^49^:

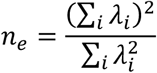

where *λ_i_* is the *i*^th^ eigenvalue of the phenotypic correlation matrix. Then, the posterior standard error of the mean was computed as

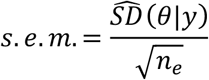

where 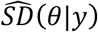 is the standard deviation of the parameter estimate across traits.

### GWAS summary statistics

We performed GWAS analyses for 26 quantitative traits and 9 common diseases in the full release of the UKB data using PLINK 1.90 (URLs). We used 348,501 unrelated individuals of European ancestry (estimated genetic relatedness from GCTA (URLs) < 0.05) and the imputed HapMap3 SNPs (URLs) provided by the UKB team^18^. We filtered SNPs with MAF<0.01, HWE test *P* value < 1×10^-6^, missing genotype rate > 0.05, or imputation info score < 0.3. We further excluded SNPs in the Human Major Histocompatibility Complex (MHC) region, resulting in a total of 1,124,198 common SNPs for the analysis. The LD correlations in the reference samples were estimated based on the effect alleles in the GWAS summary data. For quantitative traits, we standardised phenotypes to mean zero and variance one and performed rank-based inverse normal transformation (RINT) within each sex group. Prior to GWAS, we pre-adjusted phenotypes by age, sex and first 10 principal components (PCs) provided by the UKB team. For the publicly available summary statistics, we downloaded the data and matched the SNPs with those in the UKB data after excluding the strand ambiguous SNPs (i.e. A/T or C/G SNPs) in addition to the quality control procedures above. For the Neale Lab GWAS summary data (URLs), we extracted 274 quantitative traits for which the GWAS was performed based on RINT phenotypes with their analysis pipeline.

### Projection based on forward simulations

We used SLiM3^28^ to run forward simulations. A 10 Mb sequence was simulated, with a proportion of new mutations (*π_m_*) had causal effects sampled from a standard normal distribution, and the rest being neutral. The mutation rate was set to 1.65×10^-8^ per base pair per individual per generation^50^, and the recombination rate was set to 1×10^-8^. A demographic model^36^ with population bottleneck, expansion and migration was used to simulate a population undergone selection for 11,000 generations. In each generation (*t*), we computed the aggregated genotypic value of all segregating causal variants and calculated the genetic variance across individuals 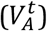. The phenotype was generated by adding a random normal deviate with a variance of 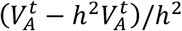 to the genetic value given a trait heritability (*h*^2^). A normalising stabilising selection model^37^ was used to link the variance-standardised phenotype (*y*) to fitness (*f*):

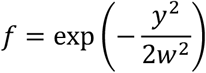

where *w*^2^ controls the decreasing rate in fitness when phenotype deviates from the optimum (at point zero), so the higher 1/*w*^2^ the stronger strength of selection. The input parameters for the simulation were mutation target size (*π_m_*), total trait heritability (*h*^2^), and strength of stabilising selection (*w*^2^). We set *π_m_*=0.025 or 0.05, *h*^2^=0.025 or 0.05, and 1/*w*^2^=0, 0.1, 0.2, 0.5, 1, 2 or 5 to obtain a map of genetic architecture of the simulated trait. In the last generation, we estimated the SNP-based heritability, polygenicity and *S* based on the common causal variants (MAF>0.01). Since we have observed the true effect size, the *S* parameter was estimated using a linear regression

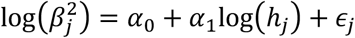

where the slope *α*_1_ is an estimate of *S* according to the BayesS model, and the residuals ***ϵ*** are independent. We repeated this simulation process 30 times and computed the mean of each genetic architecture parameter in each of the scenarios.

To facilitate the projection of real-trait genetic architecture, we first mapped the genetic architecture parameters estimated from the simulated traits in a polar coordinate, with the estimates scaled by the respective maximum values across simulation scenarios to give the same range between zero and one for each parameter. Connecting the values in the polar coordinate manifested a triangle that was unique to a simulation scenario. Similarly, we mapped the estimated genetic architecture parameters for the real traits and formed a triangle by scaling each parameter estimate by the maximum value across traits. For each real trait, we then sought to project it onto the simulation scenario that had the most similar triangle by minimising the sum of the differences in internal angles between the two triangles. Although the point estimates of the genetic architecture parameters were used to form the triangle, it can be seen that the estimation variation did not have substantial effect on the shape of the triangle (Supplementary Fig. 21). Finally, we computed the median values of the parameter estimates for each trait category, and projected them onto the map of genetic architecture using the same method.

## URLs

UK Biobank: https://www.ukbiobank.ac.uk

GERA: https://www.ncbi.nlm.nih.gov/proiects/gap/cgi-bin/study.cgi?study_id=phs000674.v2.p2

UKB GWAS summary data from the Neale Lab: http://www.nealelab.is/uk-biobank

PLINK 1.90: https://www.cog-genomics.org/plink2

SLiM3: https://messerlab.org/slim

GCTA: https://cnsgenomics.com/software/gcta

GCTB: https://cnsgenomics.com/software/gctb

HapMap3: https://www.sanger.ac.uk/resources/downloads/human/hapmap3.html

baseline-LD annotations: https://data.broadinstitute.org/alkesgroup/LDSCORE

## Acknowledgements

This research was supported by the Australian Research Council (DP160101343, DP160101056, and FT180100186), and the Australian National Health and Medical Research Council (1107258, 1078901, 1078037, and 1113400). This study makes use of data from dbGaP (accession: phs000788) and UK Biobank Resource (application number: 12505). A full list of acknowledgements for these datasets can be found in the Supplementary Note.

## Author contributions

J.Y. and J.Z. conceived the study and designed the experiment. J.Z. derived the analytical methods, conducted all analyses, and developed the software with assistance and guidance from J.Y., P.M.V., N.R.W., M.E.G., L.R.L-J., Y.W., H.W., Z.Z. and L.Y. A. X. and L. J. curated the GWAS summary statistics from published studies. K.E.K., L.Y. and Z.Z. performed the initial preparation and quality control of the UK Biobank data. J.Z. and J.Y. wrote the manuscript with the participation of all authors. All authors reviewed and approved the final manuscript.

## Data availability

This study makes use of individual-level genotype and phenotype data from UK Biobank Resource (application number: 12505) as well as GWAS summary data and functional genomic annotation data from the public domain (URLs).

## Code availability

SBayesS and SBayesS-strat have been implemented in a software tool called GCTB (genome-wide complex trait Bayesian analyses), the computer code for which is freely available at http://cnsgenomics.com/software/gctb.

