## Supplementary Note for "Bayesian analysis of GWAS summary data reveals differential signatures of natural selection across human complex traits and functional genomic categories"

#### 1 SBayesS model

Let us consider an individual-level data-based multiple regression model in a GWAS data set:

$$\mathbf{y} = \mathbf{X}\boldsymbol{\beta} + \mathbf{e} \tag{1}$$

where $\mathbf{y}$ is the vector of phenotypes adjusted for all fixed effects, $\mathbf{X}$ is the column-centered genotype matrix, $\boldsymbol{\beta}$ is the vector of SNP effects, and $\mathbf{e}$ is the vector of residuals with $Var(\mathbf{e}) = \mathbf{I}\sigma_e^2$ for a sample of unrelated individuals. Assuming Hardy-Weinberg equilibrium (HWE), the variance of the genotypes of SNP $j$ is $h_j = 2p_jq_j$, where $p_j$ is the minor allele frequency (MAF) and $q_j = 1 - p_j$. Let $\mathbf{D}$ be a diagonal matrix with $D_j = \mathbf{X}_j'\mathbf{X}_j = h_jn_j$, where $n_j$ is per-SNP sample size. Multiplying both sides of (1) by $\mathbf{D}^{-1}\mathbf{X}'$ gives

$$\mathbf{D^{-1}X'y} = \mathbf{D^{-1}X'X}\boldsymbol{\beta} + \mathbf{D^{-1}X'e} \tag{2}$$

Note that $\mathbf{D^{-1}X'y} = \mathbf{b}$, the vector of least square estimates of SNP marginal effects from GWAS, and $\mathbf{D^{-1}X'X} = \mathbf{D}^{-\frac{1}{2}}\mathbf{B}\mathbf{D}^{\frac{1}{2}}$ where $\mathbf{B} = \mathbf{D}^{-\frac{1}{2}}\mathbf{X'X}\mathbf{D}^{-\frac{1}{2}}$ is the linkage disequilibrium (LD) correlation matrix among all SNPs [1]. Let $\boldsymbol{\epsilon} = \mathbf{D^{-1}X'e}$. Then, (2) can be written as

$$\mathbf{b} = \mathbf{D}^{-\frac{1}{2}}\mathbf{B}\mathbf{D}^{\frac{1}{2}}\boldsymbol{\beta} + \boldsymbol{\epsilon} \tag{3}$$

Or, in a scalar form,

$$b_j = \sum_{k=1}^{m} \sqrt{\frac{h_kn_k}{h_jn_j}} B_{jk}\beta_k + \epsilon_j \tag{4}$$

with $m$ being the total number of SNPs. Let $\sigma_{X_j}^2$, $\sigma_{X_k}^2$ and $\sigma_{X_j,X_k}$ denote the genotype variance of SNP $j$ and $k$ and their covariance. Then, (4) can be simplified to

$$b_j = \sum_{k=1}^{m} \sqrt{\frac{n_k}{n_j}} \beta_{X_j,X_k}\beta_k + \epsilon_j \tag{5}$$

where $\beta_{X_j,X_k} = \sigma_{X_j,X_k}/\sigma_{X_j}^2$ is the regression of SNP $k$ on that of SNP $j$. In other words, we model the marginal effect of each SNP as a linear combination of other SNP effects with the weights being a function of regression coefficient of SNP genotypes and per-SNP sample sizes. In contrast to the identity structure of residual variance in

(1), the residuals in (3) are not independent in the presence of LD, because

$$\begin{aligned} Var(\boldsymbol{\epsilon}) &= Var\left(\mathbf{D^{-1}X'e}\right) \\ &= \mathbf{D^{-1}X'XD^{-1}}\sigma_e^2 \\ &= \mathbf{D^{-\frac{1}{2}}BD^{-\frac{1}{2}}}\sigma_e^2 \end{aligned} \quad (6)$$

Let $\mathbf{W} = \mathbf{D^{-\frac{1}{2}}BD^{\frac{1}{2}}}$ and $\mathbf{R} = \mathbf{D^{-\frac{1}{2}}BD^{-\frac{1}{2}}}$. Finally, from (3) we have

$$\mathbf{b} = \mathbf{W}\boldsymbol{\beta} + \boldsymbol{\epsilon} \quad (7)$$

with $Var(\boldsymbol{\epsilon}) = \mathbf{R}\sigma_e^2$. This is a generic form of summary-data-based Bayesian regressions (SBR), which is similar to the RSS model of Zhu and Stephens [2].

As in BayesS [3], we assume the effect size is related to MAF through parameter S:

$$\beta_j \begin{cases} \sim N\left(0, h_j^S \sigma_\beta^2\right), & \pi \\ = 0, & 1-\pi \end{cases} \quad (8)$$

where $S$, $\sigma_\beta^2$ and $\pi$ are considered as unknown. The prior for $S$ is a standard normal distribution

$$S \sim N(0,1)$$

The prior for $\pi$ is a uniform between zero and one, namely

$$\pi \sim Beta(1,1)$$

The prior for $\sigma_\beta^2$ is a scaled inverse chi-square distribution

$$\sigma_\beta^2 \sim \nu_\beta \tau_\beta^2 \chi_{\nu_\beta}^{-2}$$

where $\nu_\beta = 4$ and

$$\tau_\beta^2 = \frac{\nu_\beta - 2}{\nu_\beta} \frac{V_P h_0^2}{\pi_0 \sum_j h_j^{S_0+1}}$$

where $V_P$ is the phenotypic variance estimated from the summary statistics (as shown below) and $h_0^2$, $\pi_0$ and $S_0$ are the prior knowledge of SNP-based heritability, $\pi$ and $S$, resepctively. Similarly, we give a scaled inverse chi-square prior for $\sigma_e^2$ in (6)

$$\sigma_e^2 \sim \nu_e \tau_e^2 \chi_{\nu_e}^{-2}$$

where $\nu_e = 4$ and

$$\tau_e^2 = \frac{\nu_e - 2}{\nu_e} V_P \left(1 - h_0^2\right)$$

We estimate the phenotypic variance of the trait following Yang et al [1]'s approach, which is based on the stadard error of the marginal SNP effect estimate from GWAS. Because

$$SE_j^2 = \frac{\sigma_j^2}{\mathbf{X}_j'\mathbf{X}_j} = \frac{\mathbf{y}'\mathbf{y} - \mathbf{X}_j'\mathbf{X}_j b_j^2}{\mathbf{X}_j'\mathbf{X}_j}$$

where $\sigma_j^2$ is the residual variance for the GWAS model fitting SNP $j$. Rearranging gives

$$V_{P,j} = \frac{\mathbf{y}'\mathbf{y}}{n_j} = D_{jj}\left(SE_j^2 + \frac{b_j^2}{n_j}\right) \tag{9}$$

The phenotypic variance $V_P$ is then calculated as the median of $V_{P,j}$ across all SNPs [1]. Since $D_{jj} = 2p_j q_j n_j$ but the allele frequencies from the publicly available summary data are often not exact, we substitute $V_{P,j}$ by $V_P$ in (9) to reestimate $p_j$ given the input values of $SE_j$, $b_j$ and $n_j$.

We call model (7) with the above prior as "SBayesS". Specifying a different prior distribution to $\beta_j$ gives SBR form of other Bayesian alphabet models. For example, a mixture prior of normals with different variances for $\beta_j$ under SBR framework becomes SBayesR [4].

#### 2 Equivalence between individual- and summary-data-based BayesS

In the summary-data-based BayesS (SBayesS), the posterior distribution of $\boldsymbol{\beta}$ is

$$\begin{aligned}
f\left(\boldsymbol{\beta} \middle| \mathbf{b}, else\right) &\propto f\left(\mathbf{b} \middle| \boldsymbol{\beta}, else\right) f\left(\boldsymbol{\beta}\right) \\
&\propto \exp\left\{-\frac{1}{2\sigma_e^2}\left(\mathbf{b} - \mathbf{W}\boldsymbol{\beta}\right)' \mathbf{R}^{-1}\left(\mathbf{b} - \mathbf{W}\boldsymbol{\beta}\right)\right\} \exp\left\{-\frac{\boldsymbol{\beta}'\mathbf{G}^{-1}\boldsymbol{\beta}}{2\sigma_\beta^2}\right\} \\
&= \exp\left\{-\frac{1}{2\sigma_e^2}\left[\mathbf{b}'\mathbf{R}^{-1}\mathbf{b} - 2\boldsymbol{\beta}'\mathbf{W}'\mathbf{R}^{-1}\mathbf{b} + \boldsymbol{\beta}'\left(\mathbf{W}'\mathbf{R}^{-1}\mathbf{W} + \frac{\sigma_e^2}{\sigma_\beta^2}\mathbf{G}^{-1}\right)\boldsymbol{\beta}\right]\right\} \\
&\propto \exp\left\{-\frac{1}{2\sigma_e^2}\left[\boldsymbol{\beta}'\left(\mathbf{W}'\mathbf{R}^{-1}\mathbf{W} + \frac{\sigma_e^2}{\sigma_\beta^2}\mathbf{G}^{-1}\right)\boldsymbol{\beta} + 2\boldsymbol{\beta}'\mathbf{W}'\mathbf{R}^{-1}\mathbf{b}\right]\right\}
\end{aligned}$$

Note that

$$\begin{aligned}
\mathbf{W}'\mathbf{R}^{-1}\mathbf{W} &= \mathbf{D}^{\frac{1}{2}}\mathbf{B}\mathbf{D}^{-\frac{1}{2}}\mathbf{D}^{\frac{1}{2}}\mathbf{B}^{-1}\mathbf{D}^{\frac{1}{2}}\mathbf{D}^{-\frac{1}{2}}\mathbf{B}\mathbf{D}^{\frac{1}{2}} \\
&= \mathbf{D}^{\frac{1}{2}}\mathbf{B}\mathbf{D}^{\frac{1}{2}} \\
&= \mathbf{D}^{\frac{1}{2}}\mathbf{D}^{-\frac{1}{2}}\mathbf{X}'\mathbf{X}\mathbf{D}^{-\frac{1}{2}}\mathbf{D}^{\frac{1}{2}} \\
&= \mathbf{X}'\mathbf{X}
\end{aligned}$$

and

$$\begin{aligned}\mathbf{W'R^{-1}b} &= \mathbf{D^{\frac{1}{2}}BD^{-\frac{1}{2}}D^{\frac{1}{2}}B^{-1}D^{\frac{1}{2}}b} \\ &= \mathbf{Db} \\ &= \mathbf{X'y}\end{aligned}$$

Thus, the posterior distribution becomes

$$f\left(\boldsymbol{\beta}|\,\mathbf{b}, else\right) \propto \exp\left\{-\frac{1}{2\sigma_e^2}\left[\boldsymbol{\beta}'\left(\mathbf{X'X}+\frac{\sigma_e^2}{\sigma_\beta^2}\mathbf{G}^{-1}\right)\boldsymbol{\beta}+2\boldsymbol{\beta}'\mathbf{X'y}\right]\right\}$$

This is equivalent to the posterior distribution given the indivdiual genotype and phenotype data. It can be shown that the above is the kernel of a multivariate normal distribution, i.e.

$$\boldsymbol{\beta}|\,\mathbf{b}, else \sim MVN\left(\mathbf{C^{-1}r}, \mathbf{C^{-1}}\boldsymbol{\sigma}_{\mathbf{e}}^{\mathbf{2}}\right)$$

where $\mathbf{C} = \mathbf{W'R^{-1}W} + \frac{\sigma_{\mathbf{e}}^{\mathbf{2}}}{\sigma_\beta^{\mathbf{2}}}\mathbf{G^{-1}} = \mathbf{X'X} + \frac{\sigma_{\mathbf{e}}^{\mathbf{2}}}{\sigma_\beta^{\mathbf{2}}}\mathbf{G^{-1}}$ and $\mathbf{r} = \mathbf{W'R^{-1}b} = \mathbf{X'y}$. It is recognized that $\mathbf{C}$ and $\mathbf{r}$ are the left- and right-hand sides of the mixed-model equations

$$\underbrace{\left[\mathbf{W'R^{-1}W}+\mathbf{G}^{-1}\lambda\right]}_{\mathbf{C}}\boldsymbol{\beta} = \underbrace{\mathbf{W'R^{-1}b}}_{\mathbf{r}} \tag{10}$$

with $\lambda = \sigma_e^2/\sigma_\beta^2$. It can be further shown that in the Gibbs sampling, the full conditional distribution of $\beta_j$ is

$$\beta_j\left|\mathbf{b}, \boldsymbol{\beta}_{-j}, else\right. \sim N\left(\frac{r_j}{C_j}, \frac{\sigma_e^2}{C_j}\right) \tag{11}$$

where

$$\begin{aligned} r_j &= D_j b_j - \sum_{k\neq j} D_j^{\frac{1}{2}} B_{jk} D_k^{\frac{1}{2}} \beta_k \\ &= \mathbf{X}_j'\mathbf{y} - \sum_{k\neq j}\mathbf{X}_j'\mathbf{X}_k\beta_k \\ C_j &= D_j + \frac{\sigma_e^2}{h_j^S\sigma_\beta^2} = \mathbf{X}_j'\mathbf{X}_j + \frac{\sigma_e^2}{h_j^S\sigma_\beta^2}\end{aligned} \tag{12}$$

#### 3 Consequence of ignoring LD sampling variation

The consequence of using a sparse LD matrix is that it can potentially bias the mean of the full conditional distribution for $\beta_j$. Let $k$ denote a SNP in nonzero LD with the target SNP and $l$ denote a SNP that does not have significant LD with the target SNP. The $r_j$ in (12) can be written as

$$\begin{aligned} r_j &= D_j b_j - \sum_k D_j^{\frac{1}{2}} B_{jk} D_k^{\frac{1}{2}} \beta_k - \sum_l D_j^{\frac{1}{2}} B_{jl} D_l^{\frac{1}{2}} \beta_l \\ &= D_j b_j - \sum_k D_j^{\frac{1}{2}} B_{jk} D_k^{\frac{1}{2}} \beta_k - t_j \\ &= r_j^* - t_j \end{aligned} \tag{13}$$

The full conditional of $\beta_j$ in (11) becomes

$$\beta_j | \, t_j \sim N\left(\frac{r_j^* - t_j}{C_j}, \frac{\sigma_e^2}{C_j}\right)$$

When reduced LD matrix is used, $B_{jl}$ is set to be zero, we therefore completely ignore $t_j$ and do not adjust the mean of the full conditional by the effects of SNPs in very low LD. However, although individual LD is trivial, the sum of them can be substantial after multiplied by $n$ because $Var\left(\sum_l \mathbf{X}_j' \mathbf{X}_l\right) = D_j Var\left(\sum_l B_{jl}\right) \propto nm$ under the null. This may break the property of MCMC and fail the Gibbs sampling.

#### 4 Modelling LD sampling variance

Suppose the observed LD correlation between SNP $j$ and $k$ equal to the true population LD ($\rho_{jk}$) plus a deviation (tilde refers to the LD reference sample):

$$\begin{aligned} B_{jk} &= \rho_{jk} + \delta_{jk} \\ \widetilde{B}_{jk} &= \rho_{jk} + \widetilde{\delta}_{jk} \end{aligned}$$

Then, the LD correlation in the GWAS sample is

$$B_{jk} = \begin{cases} \widetilde{B}_{jk} + \left(\delta_{jk} - \widetilde{\delta}_{jk}\right) & \text{if } \rho_{jk} \neq 0 \\ \delta_{jk} & \text{if } \rho_{jk} = 0 \end{cases} \tag{14}$$

Let $\Delta_{jk}$ denote the unobserved quantity in (14), i.e.

$$\Delta_{jk} = \begin{cases} \delta_{jk} - \widetilde{\delta}_{jk} & \text{if } \rho_{jk} \neq 0 \\ \delta_{jk} & \text{if } \rho_{jk} = 0 \end{cases}$$

In (7), we can write

$$\mathbf{W} = \mathbf{W}^{+} + \mathbf{W}^{-} \tag{15}$$

where $\mathbf{W}^{+} = \mathbf{D}^{-\frac{1}{2}}\widetilde{\mathbf{B}}\mathbf{D}^{\frac{1}{2}}$ is the observed data, and $\mathbf{W}^{-} = \mathbf{D}^{-\frac{1}{2}}\boldsymbol{\Delta}\mathbf{D}^{\frac{1}{2}}$ is not observed. Substituing (15) in (7) give

$$\mathbf{b} = \mathbf{W}^{+}\boldsymbol{\beta} + \boldsymbol{\eta} \tag{16}$$

where $\boldsymbol{\eta} = \mathbf{W}^{-}\boldsymbol{\beta} + \boldsymbol{\epsilon}$ are the new residuals that contain difference in sampling deviation of LD between the GWAS and reference samples when the population LD is not zero, and sampling deviation of LD in the GWAS sample when the population LD is zero.

Conditional on $\boldsymbol{\Delta}$, the residual variance is

$$Var\left(\boldsymbol{\eta}\,|\boldsymbol{\Delta}\,\right) = \mathbf{W}^{-\prime}\mathbf{G}\mathbf{W}^{-}\sigma_{\beta}^{2} + \mathbf{R}\sigma_{e}^{2} \tag{17}$$

with $\mathbf{R} = \mathbf{D}^{-\frac{1}{2}}\widetilde{\mathbf{B}}\mathbf{D}^{-\frac{1}{2}}$.

Considering both $\boldsymbol{\beta}$ and $\boldsymbol{\Delta}$ as random, the diagonal value of $Var(\boldsymbol{\eta})$ in 16 is

$$\begin{aligned}
Var(\eta_j) &= E\left[Var(\eta_j|\,\boldsymbol{\Delta}_j)\right] + Var\left[E(\eta_j|\,\boldsymbol{\Delta}_j)\right] \\
&= E\left[\left(\mathbf{W}_j^-\right)'\mathbf{G}\mathbf{W}_j^-\sigma_\beta^2\pi + D_j^{-1}\sigma_e^2\right] + 0 \\
&= E\left[\sum_{k=1}^{m} D_j^{-1}\Delta_{jk}^2 D_k G_k \sigma_\beta^2\pi\right] + D_j^{-1}\sigma_e^2 \\
&= D_j^{-1}\sigma_\beta^2\pi E\left[D_k G_k\right] E\left[\sum_k \Delta_{jk}^2\right] + D_j^{-1}\sigma_e^2 \\
&= D_j^{-1}\sigma_\beta^2\pi \frac{\sum_k (2p_k q_k)^{S+1} n_j}{m} E\left[\sum_k \Delta_{jk}^2\right] + D_j^{-1}\sigma_e^2 \\
&= D_j^{-1}\left(\pi\sum_k (2p_k q_k)^{S+1}\sigma_\beta^2\right)\frac{n_j}{m} E\left[\sum_k \Delta_{jk}^2\right] + D_j^{-1}\sigma_e^2 \\
&= D_j^{-1}\sigma_g^2\frac{n_j}{m} E\left[\sum_k \Delta_{jk}^2\right] + D_j^{-1}\sigma_e^2 \\
&= D_j^{-1}\left(\frac{n_j}{m}\sum_k E\left[\Delta_{jk}^2\right]\sigma_g^2 + \sigma_e^2\right) \\
&= D_j^{-1}\left(\frac{n_j}{m}\sum_k Var(\Delta_{jk})\sigma_g^2 + \sigma_e^2\right) \qquad (18)
\end{aligned}$$

When $\rho_{jk} \neq 0$,

$$\begin{aligned}
Var(\Delta_{jk}) &= Var(\delta_{jk}) + Var\left(\widetilde{\delta}_{jk}\right) - 2Cov\left(\delta_{jk}, \widetilde{\delta}_{jk}\right) \\
&= \frac{\left(1-\rho_{jk}^2\right)^2}{n_j} + \frac{\left(1-\rho_{jk}^2\right)^2}{\widetilde{n}_j} - 2Cov\left(\delta_{jk}, \widetilde{\delta}_{jk}\right) \qquad (19)
\end{aligned}$$

If there is no sample overlap between the LD reference and GWAS samples, then $Cov\left(\delta_{jk}, \widetilde{\delta}_{jk}\right) = 0$. If we compute LD from the GWAS sample itself, then $Var(\Delta_{jk}) = 0$.

When $\rho_{jk} = 0$,

$$Var(\Delta_{jk}) = Var(\delta_{jk}) = \frac{1}{n_j}$$

If we estimate $\rho_{jk}$ by $\widetilde{B}_{jk}$ and assume there is no sample overlap between the LD reference and GWAS samples, substituting these results in (18) gives

$$Var(\eta_j) = D_j^{-1}\left[\left(\frac{n_j}{m}s_j^2 + \frac{m_j^0}{m}\right)\sigma_g^2 + \sigma_e^2\right] \qquad (20)$$

where $m_j^0$ is the number of SNPs not in population LD with SNP $j$, $\sigma_g^2$ is the trait genetic variance, and

$$s_j^2 = \sum_{k=1}^{m_j} \left[ \frac{\left(1-\rho_{jk}^2\right)^2}{n_j} + \frac{\left(1-\rho_{jk}^2\right)^2}{\widetilde{n}_j} - 2Cov\left(\delta_{jk}, \widetilde{\delta}_{jk}\right) \right]$$

is the total sampling variance for non-zero LD. In practice, we approximate $\rho_{jk}$ by $\widetilde{B}_{jk}$.

Similarly, considering both $\boldsymbol{\beta}$ and $\boldsymbol{\Delta}$ as random, the off-diagonal value of $Var(\boldsymbol{\eta})$ in 16 is

$$\begin{aligned}
Cov(\eta_j, \eta_k) &= E\left[Cov\left(\eta_j, \eta_k | \boldsymbol{\Delta}_j, \boldsymbol{\Delta}_k\right)\right] + Cov\left[E\left(\eta_j | \boldsymbol{\Delta}_j\right), E\left(\eta_k | \boldsymbol{\Delta}_k\right)\right] \\
&= E\left[\left(\mathbf{W}_j^-\right)' \mathbf{G}\mathbf{W}_k^- \sigma_\beta^2 + D_j^{-\frac{1}{2}} \widetilde{B}_{jk} D_k^{-\frac{1}{2}} \sigma_e^2\right] + 0 \\
&= E\left[\sum_l D_j^{-\frac{1}{2}} D_k^{-\frac{1}{2}} \Delta_{jl} \Delta_{kl} D_l G_l \sigma_\beta^2 \pi\right] + D_j^{-\frac{1}{2}} \widetilde{B}_{jk} D_k^{-\frac{1}{2}} \sigma_e^2 \\
&= 0 + D_j^{-\frac{1}{2}} \widetilde{B}_{jk} D_k^{-\frac{1}{2}} \sigma_e^2 \\
&= R_{jk} \sigma_e^2
\end{aligned}$$

Thus, the sampling variance of LD correlations only affect the diagonal but not off-diagonal values of the residual variance.

### 5 MCMC sampling scheme

The joint distribution of the data and parameters in model (7) is

$$\begin{aligned}
f\left(\mathbf{b}, \boldsymbol{\beta}, \pi, S, \sigma_\beta^2, \sigma_e^2\right) &\propto \left|\mathbf{R}\sigma_e^2\right|^{-\frac{1}{2}} \exp\left\{-\frac{1}{2\sigma_e^2} \left(\mathbf{b} - \mathbf{W}\boldsymbol{\beta}\right)' \mathbf{R}^{-1} \left(\mathbf{b} - \mathbf{W}\boldsymbol{\beta}\right)'\right\} \\
&\times \prod_{j=1}^m \left[\left(h_j^S \sigma_\beta^2\right)^{-\frac{1}{2}} \exp\left\{-\frac{\beta_j^2}{2h_j^S \sigma_\beta^2}\right\} \pi + \phi(1-\pi)\right] \\
&\times \exp\left\{-\frac{S^2}{2}\right\} \\
&\times \left(\sigma_\beta^2\right)^{-\frac{2+\nu_\beta}{2}} \exp\left\{-\frac{\nu_\beta \tau_\beta^2}{2\sigma_\beta^2}\right\} \\
&\times \left(\sigma_e^2\right)^{-\frac{2+\nu_e}{2}} \exp\left\{-\frac{\nu_e \tau_e^2}{2\sigma_e^2}\right\}
\end{aligned}$$

To obtain a joint posterior sample for parameter inference, we iteratively sample each parameter from its full conditional distribution. Except $S$, the full conditional distribution has a closed form for all the parameters, as shown below.

To deal with the mixture prior for $\beta_j$, we introduce an indicator variable $\delta_j$

$$\delta_j \sim Bernoulli(\pi)$$

such that

$$\beta_j \begin{cases} \sim N(0, h_j^S \sigma_\beta^2), & \delta_j = 1 \\ = 0, & \delta_j = 0 \end{cases}$$

We first sample $\delta_j$ unconditional on $\beta_j$ and then sample $\beta_j$ conditional on $\delta_j$, which has been shown to have slightly better mixing. The full conditional distribution for $\delta_j$ is

$$\delta_j \left| \mathbf{b}, else \right. \sim Bernoulli(\hat{\pi})$$

where

$$\begin{aligned} \hat{\pi} &= \frac{f\left(\mathbf{b} \left| \delta_j = 1, else \right.\right)\pi}{f\left(\mathbf{b} \left| \delta_j = 1, else \right.\right)\pi + f\left(\mathbf{b} \left| \delta_j = 0, else \right.\right)(1-\pi)} \\ &= \left[1 + \frac{f\left(\mathbf{b} \left| \delta_j = 0, else \right.\right)}{f\left(\mathbf{b} \left| \delta_j = 1, else \right.\right)} \frac{1-\pi}{\pi}\right]^{-1} \end{aligned} \tag{21}$$

It is obvious that

$$f\left(\mathbf{b} \left| \delta_j = 0, else \right.\right) = \left(2\pi \left|\mathbf{R}\right| \sigma_e^2\right)^{-\frac{1}{2}} \exp\left\{-\frac{1}{2\sigma_e^2} \mathbf{b}_{adj}' \mathbf{R}^{-1} \mathbf{b}_{adj}\right\} \tag{22}$$

where $\mathbf{b}_{adj} = \mathbf{b} - \sum_{k \neq j} \mathbf{W}_k \beta_k$ is the adjusted $\mathbf{b}$ for all the other SNP effects except SNP $j$. Fortunately, we do not need to compute this quantity because it will be cancelled in the likelihood ratio in (21), as shown below. For $f\left(\mathbf{b} \left| \delta_j = 1, else \right.\right)$, to be unconditional on $\beta_j$, we compute

$$\begin{aligned} f\left(\mathbf{b} \left| \delta_j = 1, else \right.\right) &= \int f\left(\mathbf{b} \left| \delta_j = 1, \beta_j, else \right.\right) f\left(\beta_j\right) d\beta_j \\ &= \left(2\pi \left|\mathbf{R}\right| \sigma_e^2\right)^{-\frac{1}{2}} \exp\left\{-\frac{1}{2\sigma_e^2} \mathbf{b}_{adj}' \mathbf{R}^{-1} \mathbf{b}_{adj}\right\} \left(\frac{\sigma_{e_j}^{2*}}{h_j^S \sigma_\beta^2 C_j^*}\right)^{\frac{1}{2}} \exp\left\{\frac{\left(r_j^*\right)^2}{2C_j^* \sigma_{e_j}^{2*}}\right\} \end{aligned} \tag{23}$$

This equation is derived based on integrating $\beta_j$ out of the joint distribution of $\mathbf{b}$ and $\beta_j$, which is closely related to the full conditional distribution of $\beta_j$, as shown next. Also see below for the definition of $\sigma_{e_j}^{2*}$, $C_j^*$ and $r_j^*$. Substituting (22) and (23) into (21) gives the full conditional probability for $\delta_j = 1$.

Let $\sigma_{e_j}^{2*} = \left(\frac{n_j}{m} s_j^2 + \frac{m_j^0}{m}\right) \sigma_g^2 + \sigma_e^2$, which explicitly models the sampling variation of LD as described above. The full conditional distribution of $\beta_j$ (11) is

$$\beta_j \left| \mathbf{b}, \boldsymbol{\beta}_{-j}, else \right. \sim N\left(\frac{r_j^*}{C_j^*}, \frac{\sigma_{e_j}^{2*}}{C_j^*}\right) \tag{24}$$

with $r_j^*$ as in (13) (ignores $t_j$ the cumulative effects of SNPs in chance LD) and $C_j^* = D_j + \frac{\sigma_{e_j}^{2*}}{h_j^S \sigma_\beta^2}$. It can be seen that instead of adjusting for the "leftover" effect from the mean, we shrink the mean towards zero while increase the variance (uncertainty) of the posterior distribution, because $\sigma_{e_j}^{2*}/C_j^* = 1/\left(D_j/\sigma_{e_j}^{2*} + 1/h_j^S \sigma_\beta^2\right)$.

Given the sampled values of $\boldsymbol{\delta}$ and $\boldsymbol{\beta}$, the full conditional distribution for $\pi$, $\sigma_\beta^2$ and $S$ is the same as in BayesS, with the sampling procedure elaborated in the Supplementary Note of Zeng et al [3].

The full conditional distribution for the residual variance $\sigma_e^2$ is

$$\sigma_e^2 \left|\mathbf{b}, else\right. \sim \nu_e \tau_e^2 \chi_{\nu_e}^{-2} \tag{25}$$

where $\nu_e = \bar{n} + \nu_{e_0}$ and $\tau_e^2 = \left(\mathbf{e}'\mathbf{e} + \nu_{e_0}\tau_{e_0}^2\right)/\nu_e$ with $\nu_{e_0}$ and $\tau_{e_0}^2$ being the prior values. The residual sum of squares $(\mathbf{e}'\mathbf{e})$ can be computed as

$$\begin{aligned}
\mathbf{e}'\mathbf{e} &= (\mathbf{y} - \mathbf{X}\boldsymbol{\beta})'(\mathbf{y} - \mathbf{X}\boldsymbol{\beta}) \\
&= \mathbf{y}'\mathbf{y} - 2\boldsymbol{\beta}'\mathbf{X}'\mathbf{y} + \boldsymbol{\beta}'\mathbf{X}'\mathbf{X}\boldsymbol{\beta} \\
&= \mathbf{y}'\mathbf{y} - 2\boldsymbol{\beta}'\mathbf{r} + \boldsymbol{\beta}'(\mathbf{r} - \mathbf{r}_{adj}) \\
&= \mathbf{y}'\mathbf{y} - \boldsymbol{\beta}'\mathbf{r} - \boldsymbol{\beta}'\mathbf{r}_{adj}
\end{aligned} \tag{26}$$

where $\mathbf{r} = \mathbf{D}\mathbf{b}$ and $\mathbf{r}_{adj} = \mathbf{r} - \mathbf{X}'\mathbf{X}\boldsymbol{\beta}$ is the adjusted right-hand-side from the right-hand-side updating strategy (see below). The total sum of squares $\mathbf{y}'\mathbf{y}$ is computed from (9) and the median is used as the estimate of $\mathbf{y}'\mathbf{y}$ in (26).

#### 6 The right-hand-side updating strategy and parallel computing

The MCMC implementation requires a computation of $r_j$ in (12) for $m \times t$ times where $m$ is the number of SNPs and $t$ is the number of MCMC iterations. This is where the most majority computing time is spent. It can be seen from (12) that the summary-data level model is already much more efficient than the individual-data level model, because the vector-by-vector products $\mathbf{X}_j'\mathbf{y}$ and $\mathbf{X}_j'\mathbf{X}_k$ are replaced by scalar products $D_j b_j$ and $D_j^{\frac{1}{2}} B_{jk} D_k^{\frac{1}{2}}$. However, the adjustment of $D_j b_j$ (the right-hand-side $\mathbf{r}$ of the mixed-model equations (10)) for all the other SNP effects is still too computationally intense given over a million of SNPs. To improve computational efficiency, we adopted the so-called right-hand-side updating algorithm for genomic prediction in the context of animal breeding [5]. We set out to compute a vector of adjusted right-hand-side

$$\mathbf{r}_{adj} = \mathbf{r} - \mathbf{C}\boldsymbol{\beta}$$

, where $\mathbf{r} = \mathbf{D}\mathbf{b}$ and $\mathbf{C} = \mathbf{D}^{\frac{1}{2}}\mathbf{B}\mathbf{D}^{\frac{1}{2}}$ (note that here $\mathbf{C}$ is defined different from that in (10)). For each SNP, we compute

$$r_j = r_{adj,j} + C_{jj}\beta_j$$

and use $r_j$ in the full conditional distribution of $\beta_j$ (24). After a new value of $\beta_j$ is sampled, we update the adjusted right-hand-side

$$\mathbf{r}_{adj}^{new} = \mathbf{r}_{adj}^{old} + \mathbf{C}_j \left(\beta_j^{old} - \beta_j^{new}\right)$$

The benefit of this updating strategy comes from three sides. First, the computation of $r_j$ for each SNP becomes trivial (reducing from vector to scalar operation). Second, the vector of $\mathbf{r}_{adj}$ needs to be updated only when either $\beta_j^{old}$ or $\beta_j^{new}$ is not zero, thus the sparse genetic architecture will lead to a substantial gain in speed. Third, due to the use of a sparse LD matrix, the vector of $\mathbf{C}_j$ has a large proportion of zero and therefore only a small fraction of $\mathbf{r}_{adj}$ according to nonzero $\mathbf{C}_j$ elements needs to be updated. In summary, the right-hand-side updating strategy substantially improves computational efficiency by taking the advantages of the sparse genetic architecture and the sparse LD correlation matrix. We further improve the efficiency by implementing a parallel computing for sampling $\beta_j$ of SNPs located on different chromosomes, and then combine results across threads to estimate the global parameters such as $\pi$, $\sigma_\beta^2$, $S$ and etc. This led to about 4 times faster when 4 cores were used with OpenMP library in our real trait analysis.

### 7 Acknowledgements

**UKB:** This study has been conducted using UK Biobank resource under Application Number 12505. UK Biobank was established by the Wellcome Trust medical charity, Medical Research Council, Department of Health, Scottish Government and the Northwest Regional Development Agency. It has also had funding from the Welsh Assembly Government, British Heart Foundation and Diabetes UK.

**GERA:** The Genetic Epidemiology Research on Adult Health and Aging study was supported by grant RC2 AG036607 from the National Institutes of Health, grants from the Robert Wood Johnson Foundation, the Ellison Medical Foundation, the Wayne and Gladys Valley Foundation and Kaiser Permanente. The authors thank the Kaiser Permanente Medical Care Plan, Northern California Region (KPNC) members who have generously agreed to participate in the Kaiser Permanente Research Program on Genes, Environment and Health (RPGEH).

### References

[1] Jian Yang, Teresa Ferreira, Andrew P Morris, Sarah E Medland, Genetic of Consortium, DIAbetes Consortium, Pamela AF Madden, Andrew C Heath, Nicholas G Martin, Grant W Montgomery, Michael N Weedon, Ruth J

Loos, Timothy M Frayling, McCarthy, Mark I, Joel N Hirschhorn, Michael E Goddard, Peter M Visscher. Conditional and joint multiple-SNP analysis of GWAS summary statistics identifies additional variants influencing complex traits. *Nat Genet*, 44(4):369, 2012.

[2] Xiang Zhu, Matthew Stephens. Bayesian large-scale multiple regression with summary statistics from genome-wide association studies. *Ann Appl Statistics*, 11(3):1561–1592, 2017.

[3] Jian Zeng, Ronald Vlaming, Yang Wu, Matthew R Robinson, Lloyd-Jones, Luke R, Loic Yengo, Chloe X Yap, Angli Xue, Julia Sidorenko, McRae, Allan F, Joseph E Powell, Grant W Montgomery, Andres Metspalu, Tonu Esko, Greg Gibson, Naomi R Wray, Peter M Visscher, Jian Yang. Signatures of negative selection in the genetic architecture of human complex traits. *Nat Genet*, strona 1, 2018.

[4] Lloyd-Jones, Luke R, Jian Zeng, Julia Sidorenko, Loic Yengo, Gerhard Moser, Kathryn E Kemper, Huanwei Wang, Zhili Zheng, Reedik Magi, Tonu Esko, Andres Metspalu, Naomi R Wray, Michael E Goddard, Jian Yang, Peter M Visscher. Improved polygenic prediction by bayesian multiple regression on summary statistics. *Biorxiv*, strona 522961, 2019.

[5] Mario PL Calus. Right-hand-side updating for fast computing of genomic breeding values. *Genet Sel Evol*, 46(1):1–11, 2014.
