## Supplementary Figures for "Bayesian analysis of GWAS summary data reveals differential signatures of natural selection across human complex traits and functional genomic categories"

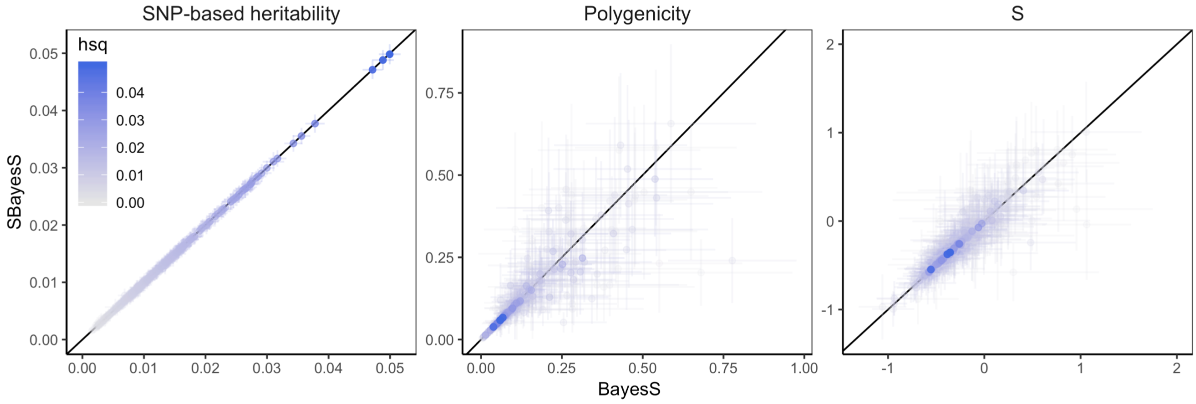


**Supplementary Figure 1** Benchmarking SBayesS with BayesS using the same data in the chromosome-wide analysis across 18 UKB traits. The comparison was based on the unrelated individuals of European ancestry in the interim release of the UKB data (max n=120k) and ~500k array genotyped common SNPs (MAF>0.01). In the SBayesS analysis, the full LD matrix that included all pairwise LD was used for each chromosome.

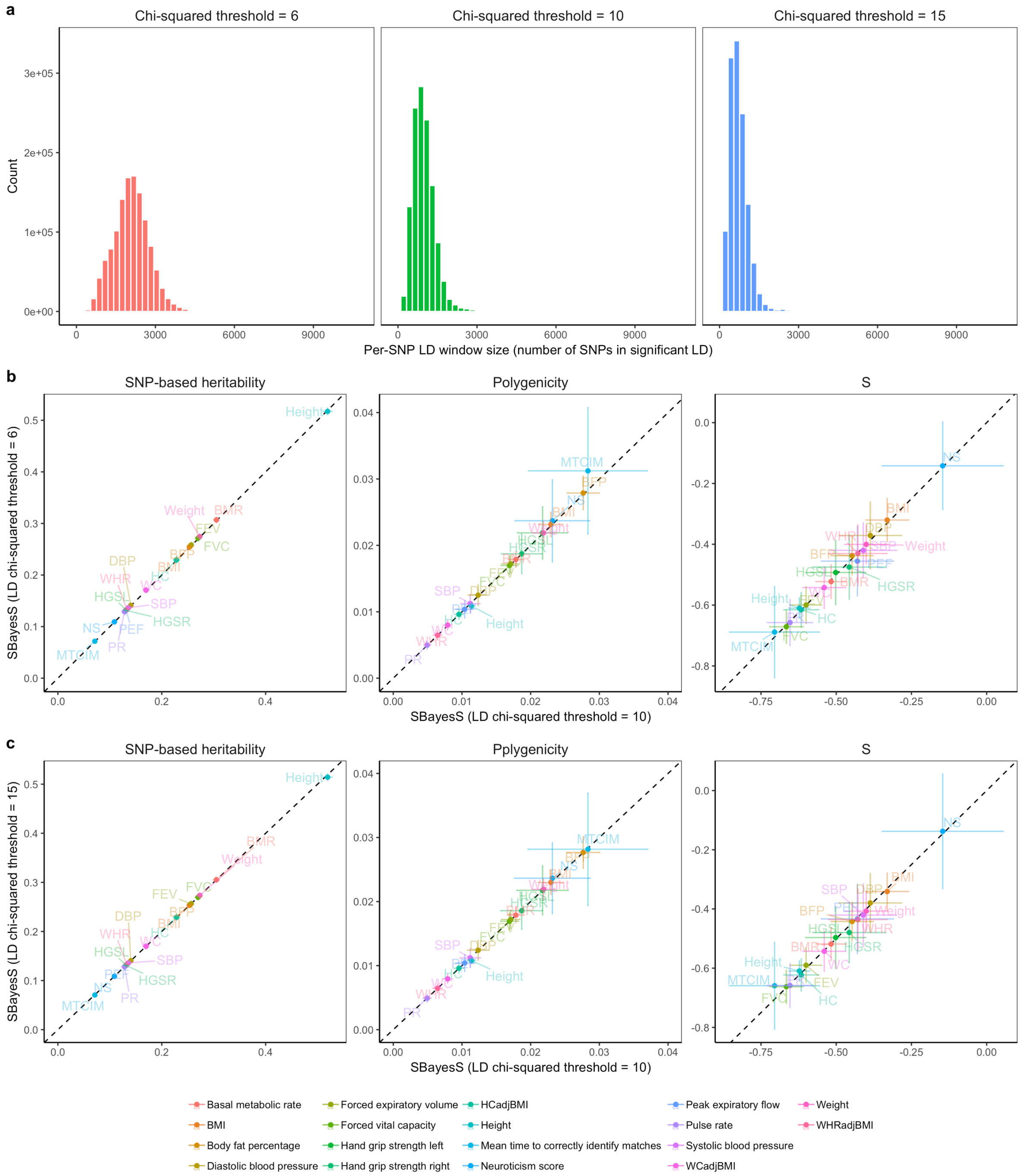


**Supplementary Figure 2** Assessing the performance of SBayesS with different chi-squared thresholds used to make the sparse LD matrix. We computed summary statistics for 18 traits using the interim release of the UKB data (max n=120k) with ~1.1 million HapMap3 common SNPs. The sparse LD matrix was computed from a random sample of 50k unrelated individuals in the full UKB data with a chi-squared threshold of 6, 10 or 15 (corresponding to a $r^2$ threshold of 1, 2 or $3\times10^{-4}$, respectively). a) Distributions of the numbers of SNPs detected in LD with the target SNP given different chi-squared thresholds; b) Comparison between SBayesS results with chi-squared thresholds of 6 and 10; c) Comparison between SBayesS results with chi-squared thresholds of 10 and 15.

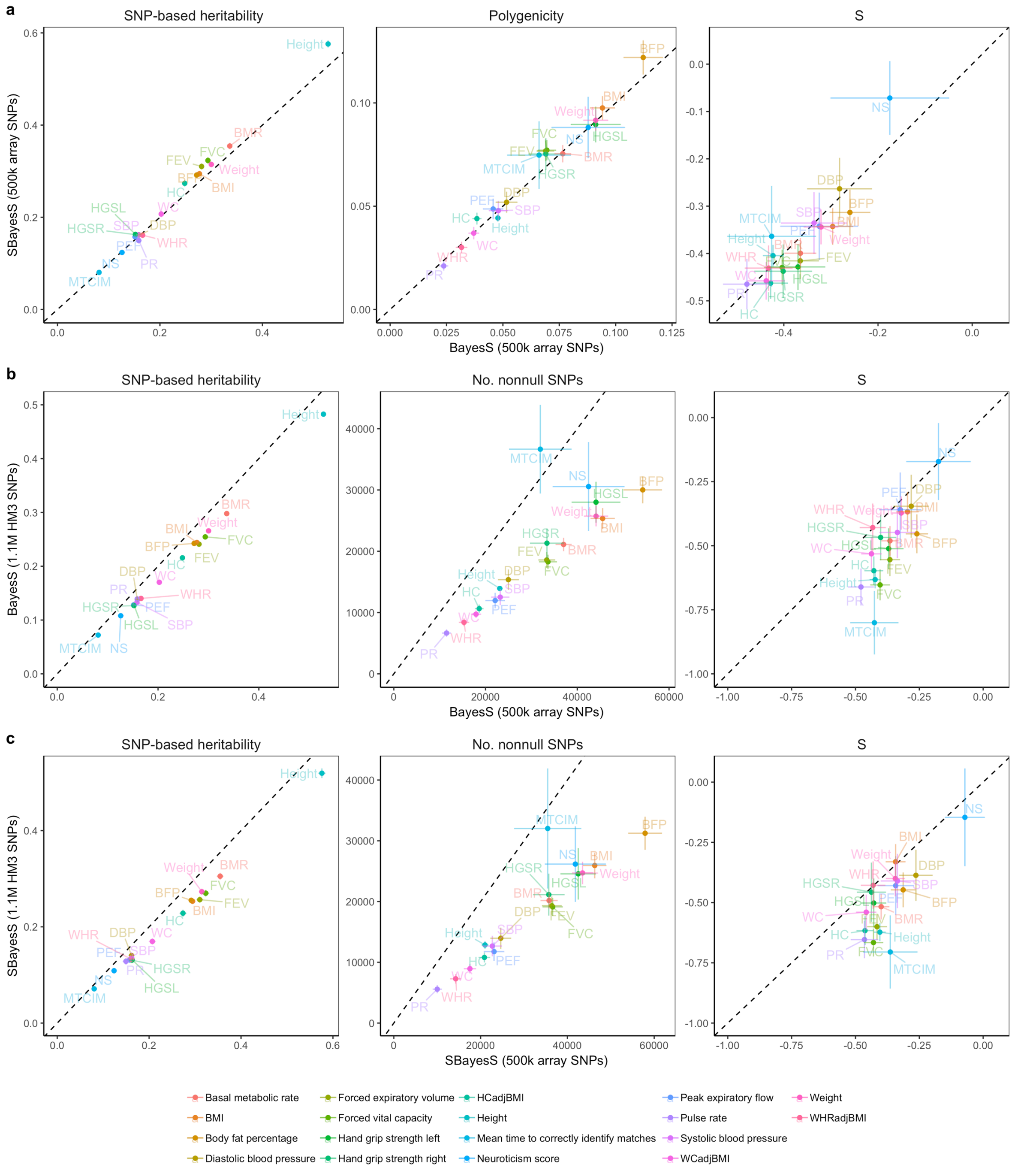


**Supplementary Figure 3** Benchmarking SBayesS with BayesS given different SNP panels. We used the unrelated individuals of European ancestry in the interim release of the UKB data (max n=120k) and two SNP panels (~500k Affymetrix array SNPs and ~1.1 million HapMap3 SNPs) for the SBayesS analysis. The sparse LD matrix was computed from a random sample of 50k unrelated individuals from the full UKB cohort at a chi-squared threshold of 10. a) Comparison between SBayesS and BayesS using array SNPs; b) Comparison between BayesS results using HapMap3 and array SNPs; c) Comparison between SBayesS results using HapMap3 and array SNPs. For a fair comparison of π between panels, the number of SNPs with nonzero effects is shown in b) and c).

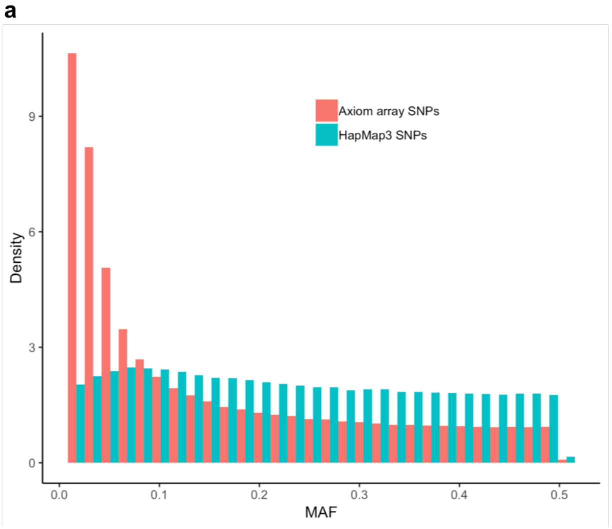


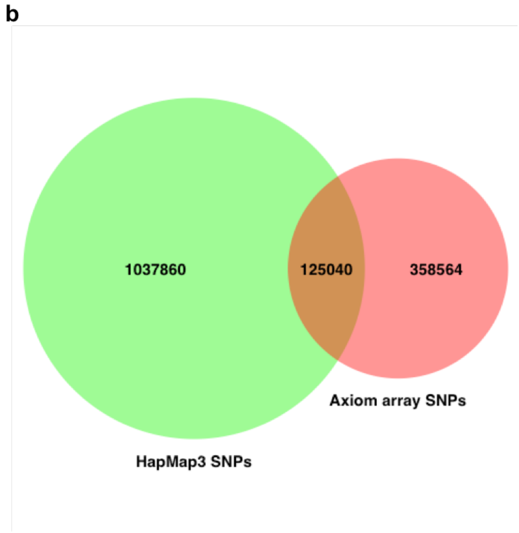


**Supplementary Figure 4** Distributions of MAF of common SNPs (MAF>1%) in Affymetrix Axiom array and HapMap3 from the interim release of the UKB data. The array SNP panel was more enriched with low-frequency SNPs and had only a small overlap with the HapMap3 SNP panel. This may explain the differences in genetic architecture parameter estimates between array and HapMap3 SNPs (Supplementary Fig. 3). For example, the array SNPs might be more efficient to capture the low-frequency variance and therefore had slightly higher total SNP-based heritability. However, the majority of the common SNPs in HapMap3 Panel is believed to have better tagging to the causal variants. Thus, it is reasonable that the polygenicity estimates were lower than those with array SNPs because the model does not need multiple SNPs in low LD with the causal variants to jointly capture the causal effects. Similarly, a stronger estimate of $S$ is expected because if the causal effects spread on multiple SNPs in LD, each SNP would have a relatively small effect size, which will dilute the signal for the relationship between effect size and MAF.

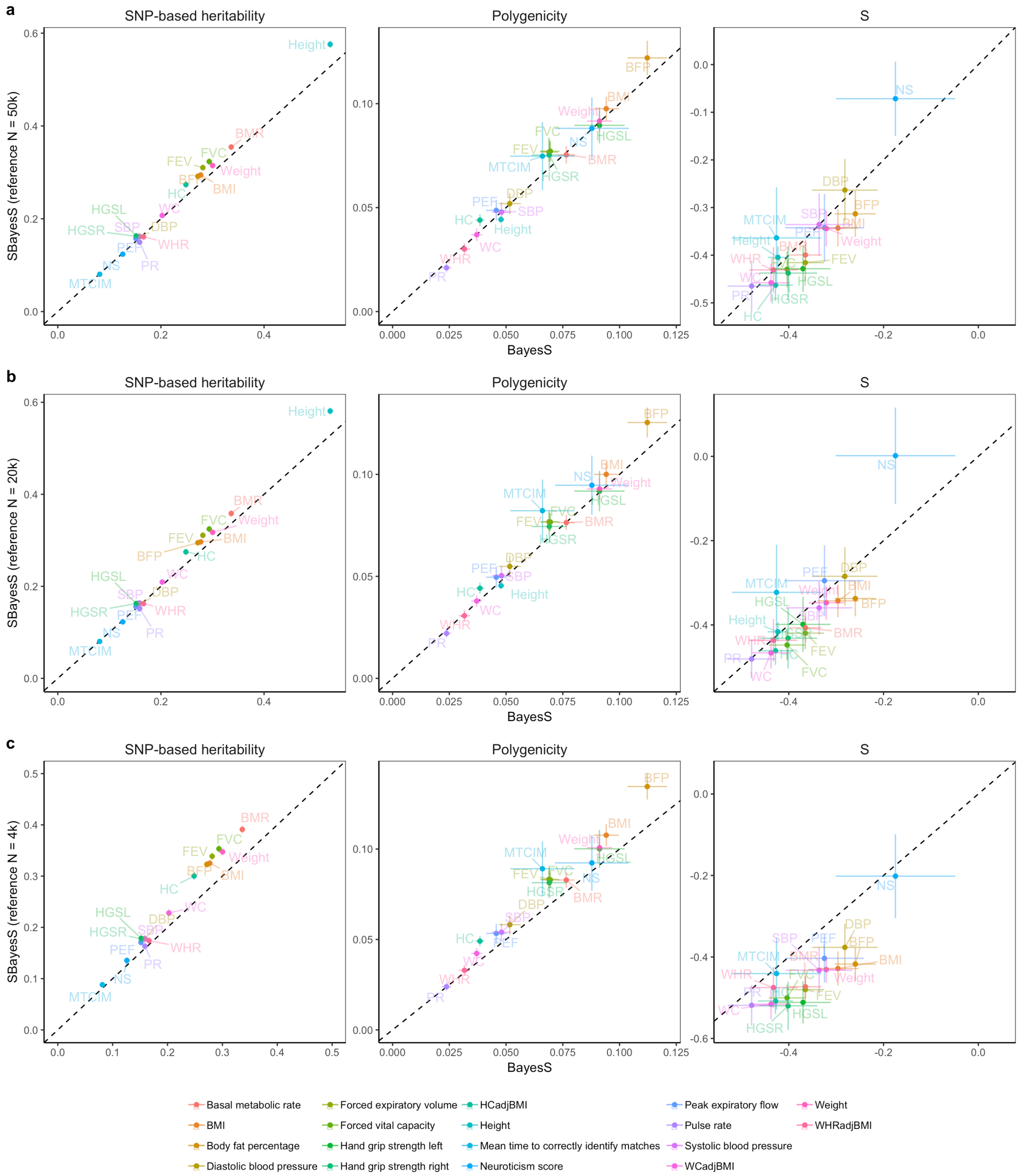


**Supplementary Figure 5** Benchmarking SBayesS with BayesS given different reference sample sizes. We used the unrelated individuals of European ancestry in the interim release of the UKB data (max n=120k) and ~500k Affymetrix array common SNPs. The sparse LD matrix was computed from a random sample of a) 50k, b) 20k or c) 4k unrelated individuals from the full UKB data at a chi-square threshold of 10. The inflation in parameter estimation increased when the reference sample size was too small. When the reference sample size was 4k, SBayesS analysis for height did not converge.

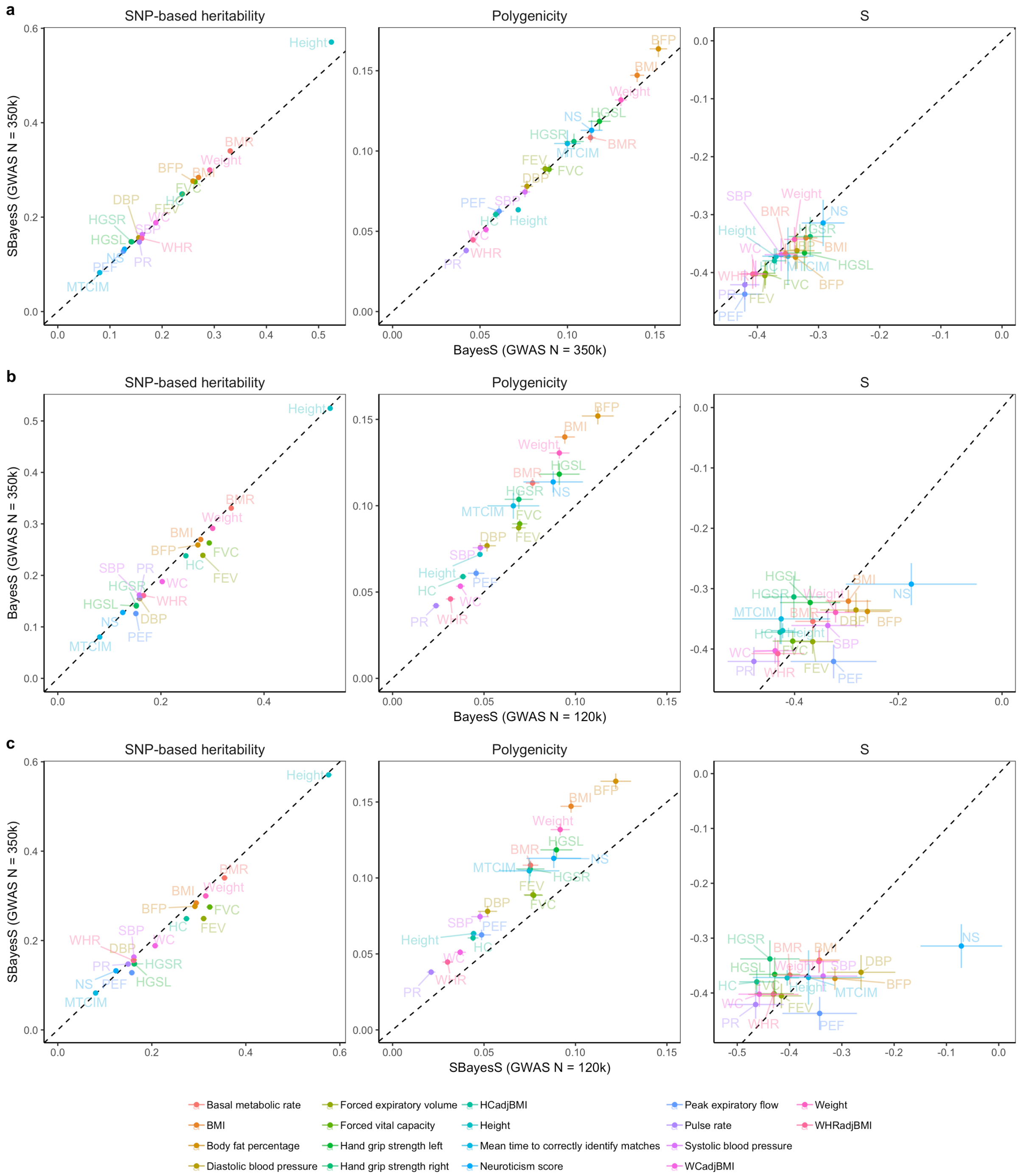


**Supplementary Figure 6** Benchmarking SBayesS with BayesS given different GWAS sample sizes. We used the unrelated individuals of European ancestry in the interim (max n=120k) and full (max n=350k) release of the UKB data ~500k Affymetrix array SNPs. The sparse LD matrix used in SBayesS was computed from a random sample of 50k unrelated individuals from the full UKB data at a chi-squared threshold of 10. a) Comparison between SBayesS and BayesS when the GWAS sample size was 350k; b) Comparison between BayesS results given GWAS sample size of 350k and 120k; c) Comparison between SBayesS results given GWAS sample size of 350k and 120k.

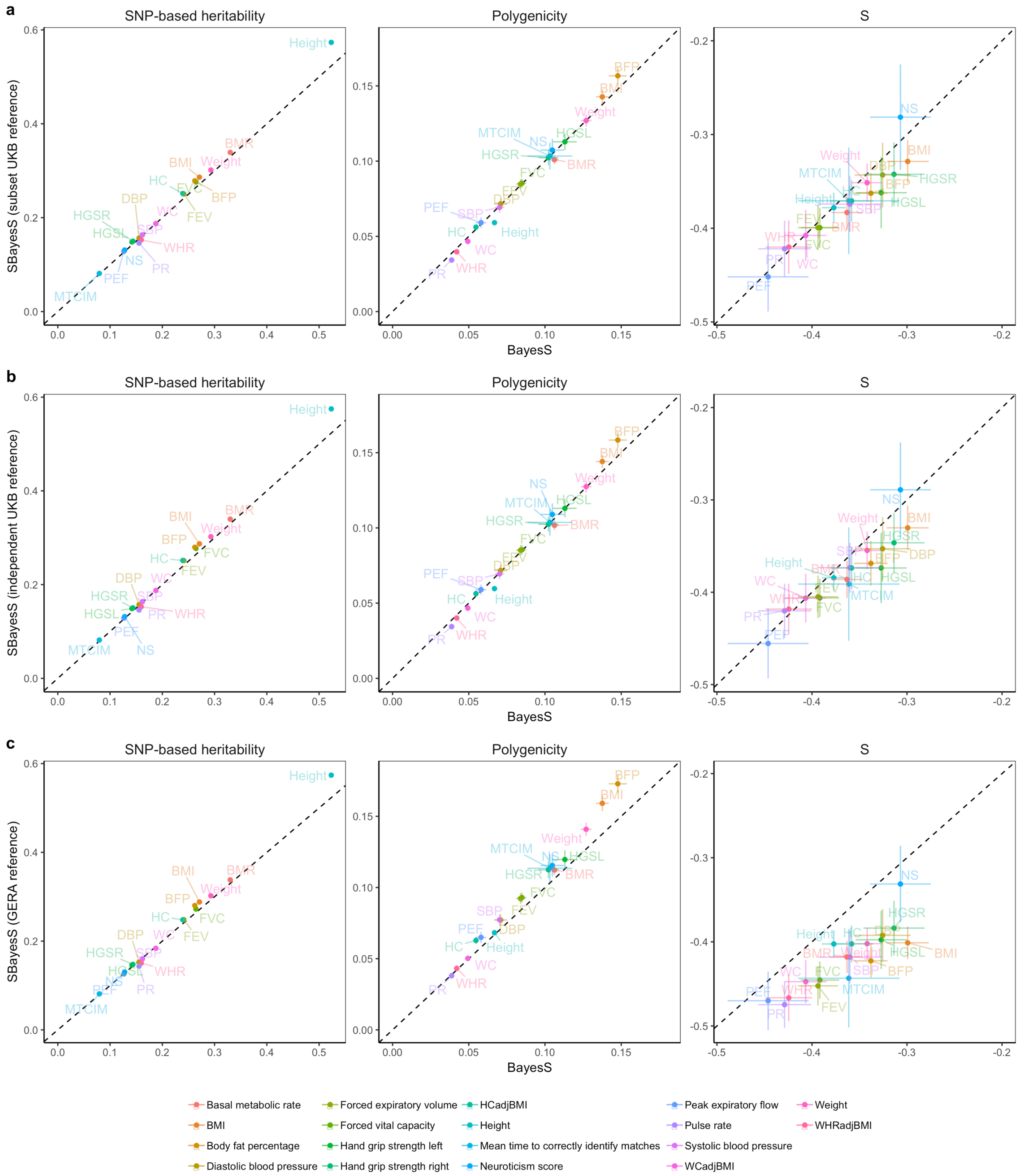


**Supplementary Figure 7** Benchmarking SBayesS with BayesS given different LD references. We used phenotypes from a random sample of 300k unrelated individuals of European ancestry from the full UKB data and ~500k Affymetrix array SNPs. The sparse LD matrix with a chi-squared threshold of 10 used in SBayesS was computed from a) a subset sample of 50k UKB individuals, b) an independent sample of 50k UKB individuals, or c) 50k unrelated individuals from the GERA dataset.

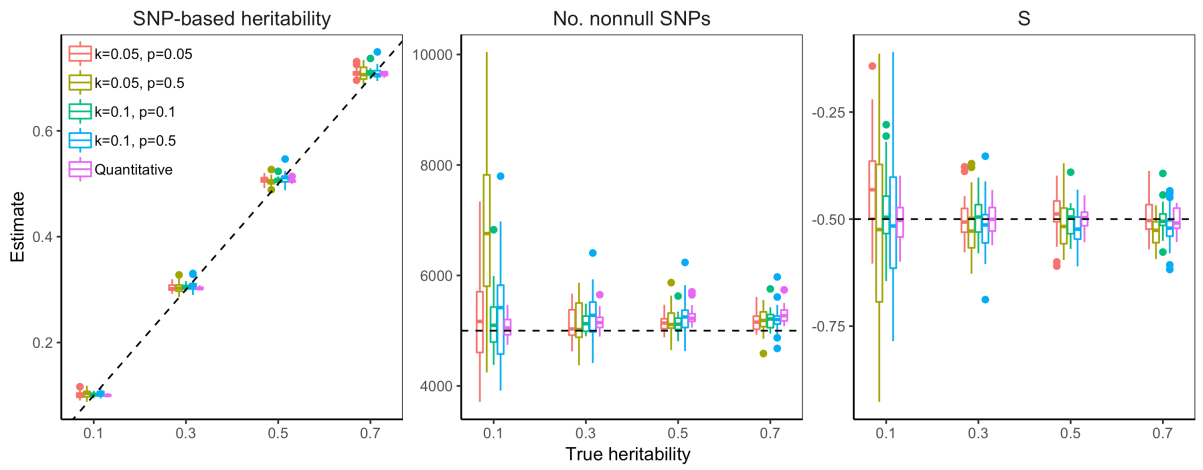


**Supplementary Figure 8** Estimation of the three genetic architecture parameters using SBayesS with simulated data for quantitative traits or case-control studies with different population (k) and sample (p) prevalence. The band inside the box is the median, the bottom and top of the box are the first and third quartiles, respectively (Q1 and Q3), and the lower and upper whiskers are Q1 – 1.5 IQR and Q3 + 1.5 IQR, respectively, where IQR = Q3 – Q1. We used the full UKB data (n=350k) for simulations, where 5k SNPs were randomly chosen from ~1.1 million HapMap3 common SNPs as causal variants with true *S*=-0.5, and the trait heritability was set to be 0.1, 0.3, 0.5 or 0.7 (at the liability scale for the binary trait). The sparse LD matrix used in SBayesS was computed from a subset sample of 50k UKB individuals with a chi-squared threshold of 10. The simulation was repeated 30 times for each scenario. When k=0.05, p=0.5 and true heritability=0.1, the polygenicity estimate tend to bias upward with large estimation variation, which is likely due to insufficient power to distinguish the model that fits only causal variants from that fits multiple SNPs in LD with the causal variants.

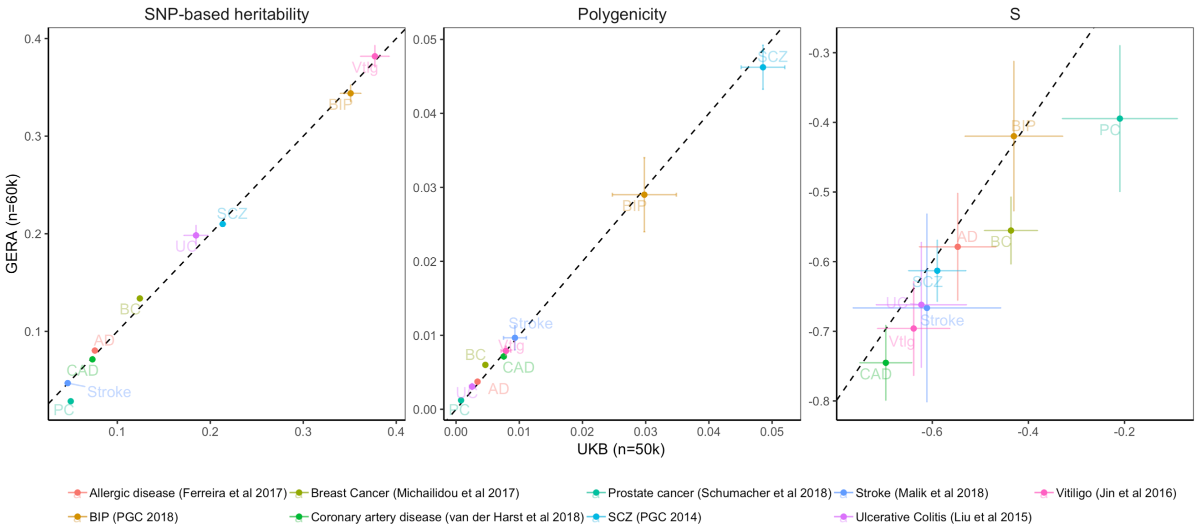


**Supplementary Figure 9** Comparison between SBayesS results based on LD from GERA and UKB (a random subsample of 50k unrelated individuals) using published GWAS summary data for 9 diseases. Each bar shows the posterior standard error of the estimate. Colours with acronyms indicate different traits, whose full names are shown at the bottom of the figure.

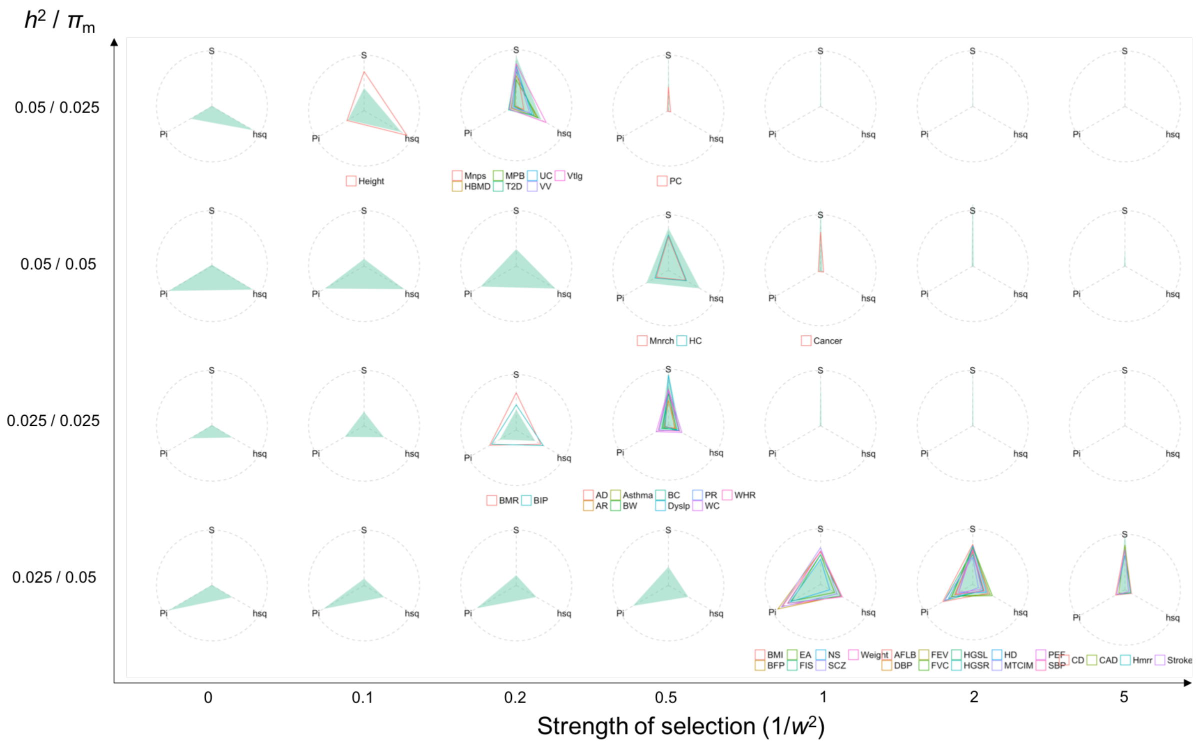


**Supplementary Figure 10** Projection of real trait genetic architecture onto the patterns observed from forward simulations with a constant effective population size. Both x and y axes are input values in the simulation, where x-axis is the strength of selection denoted as one over the variance ($w^2$) of the phenotypes surrounding at the optimum fitness value (a classic model for stabilizing selection), and y-axis is the ratio of the heritability at all causal variants ($h^2$) over the proportion of mutational targets ($\pi_m$), which is proportional to the per-mutation heritability. Each circle consists of the values of SNP-based heritability, polygenicity and S, which are scaled by the maximum values across simulated or real traits such that the three parameters have the same scale from zero to one. The green shadow indicates the computed values for the three genetic architecture parameters at the common causal variants (MAF>0.01) in the last generation of the forward simulation given the input values at x and y axes. The hollow triangle shows the estimated genetic architecture for a UKB trait. The projection of UKB trait to the map from simulation was done by minimizing the sum of the difference in each of the three angles. The forward simulation was based on a 10 Mb sequence and repeated 30 times in each scenario. A constant effective population size of 10,000 was used in the simulation.

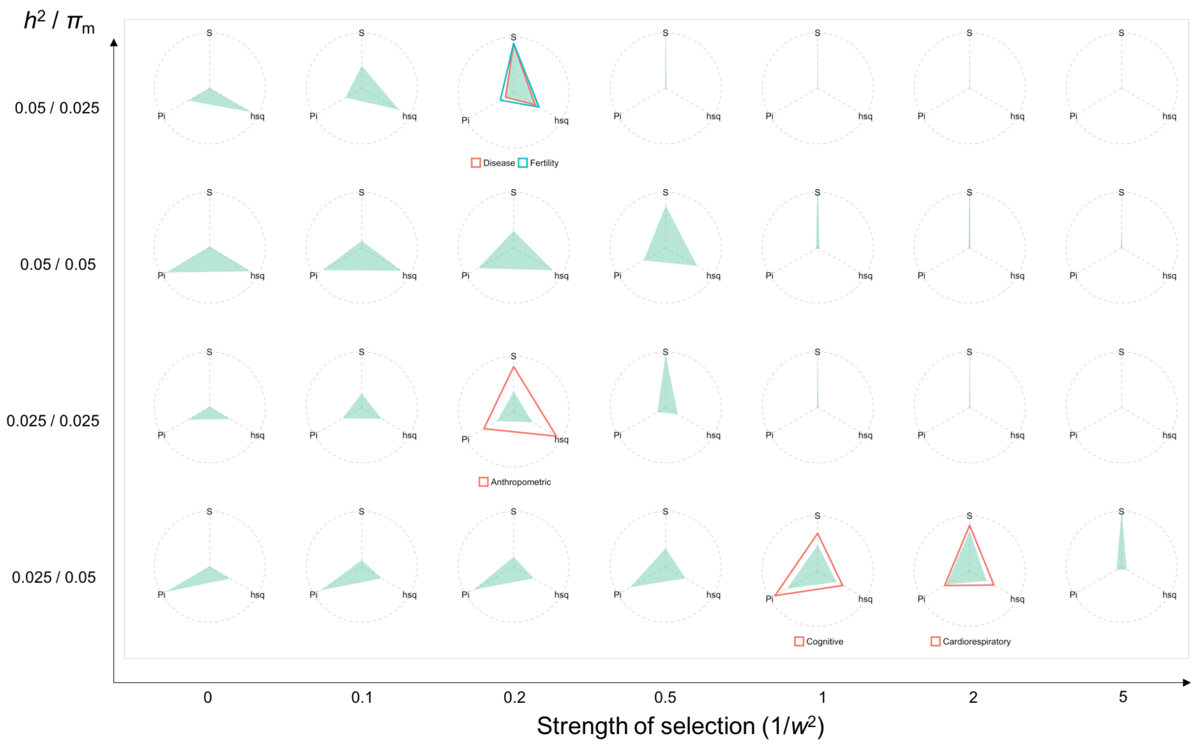


**Supplementary Figure 11** Projection of the observed genetic architecture averaged over each trait category onto the patterns observed from forward simulations with a constant effective population size. Both x and y axes are input values in the simulation, where x-axis is the strength of selection denoted as one over the variance ($w^2$) of the phenotypes surrounding at the optimum fitness value (a classic model for stabilizing selection), and y-axis is the ratio of the heritability at all causal variants ($h^2$) over the proportion of mutational targets ($\pi_m$), which is proportional to the per-mutation heritability. Each circle consists of the values of SNP-based heritability, polygenicity and S, which are scaled by the maximum values across simulated or real traits such that the three parameters have the same scale from zero to one. The green shadow indicates the computed values for the three genetic architecture parameters at the common causal variants (MAF>0.01) in the last generation of the forward simulation given the input values at x and y axes. The hollow triangle shows the estimated genetic architecture for each trait category. The projection of the observed genetic architecture to the map from simulation was done by minimizing the sum of the difference in each of the three angles. The forward simulation was based on a 10 Mb sequence and repeated 30 times in each scenario. A constant effective population size of 10,000 was used in the simulation.

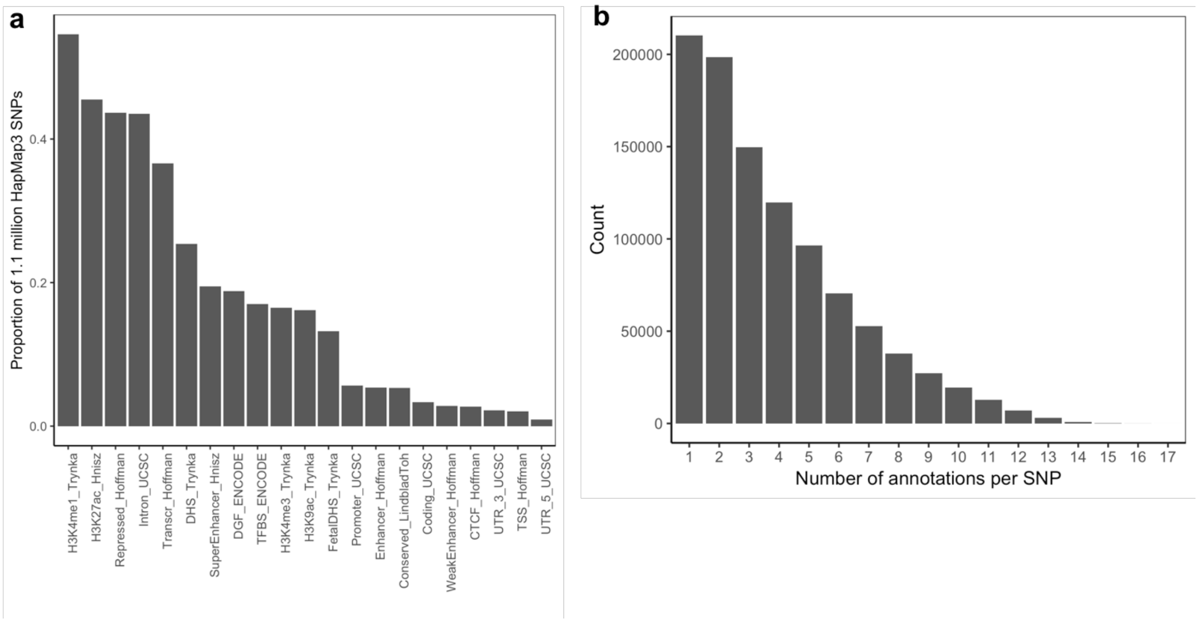


**Supplementary Figure 12** Summary of the 21 functional annotation categories from the LDSC baseline model. a) The proportion of 1.1 million HapMap3 common SNPs used in the analysis in each functional category. b) The distribution of the number of annotations for each SNP.

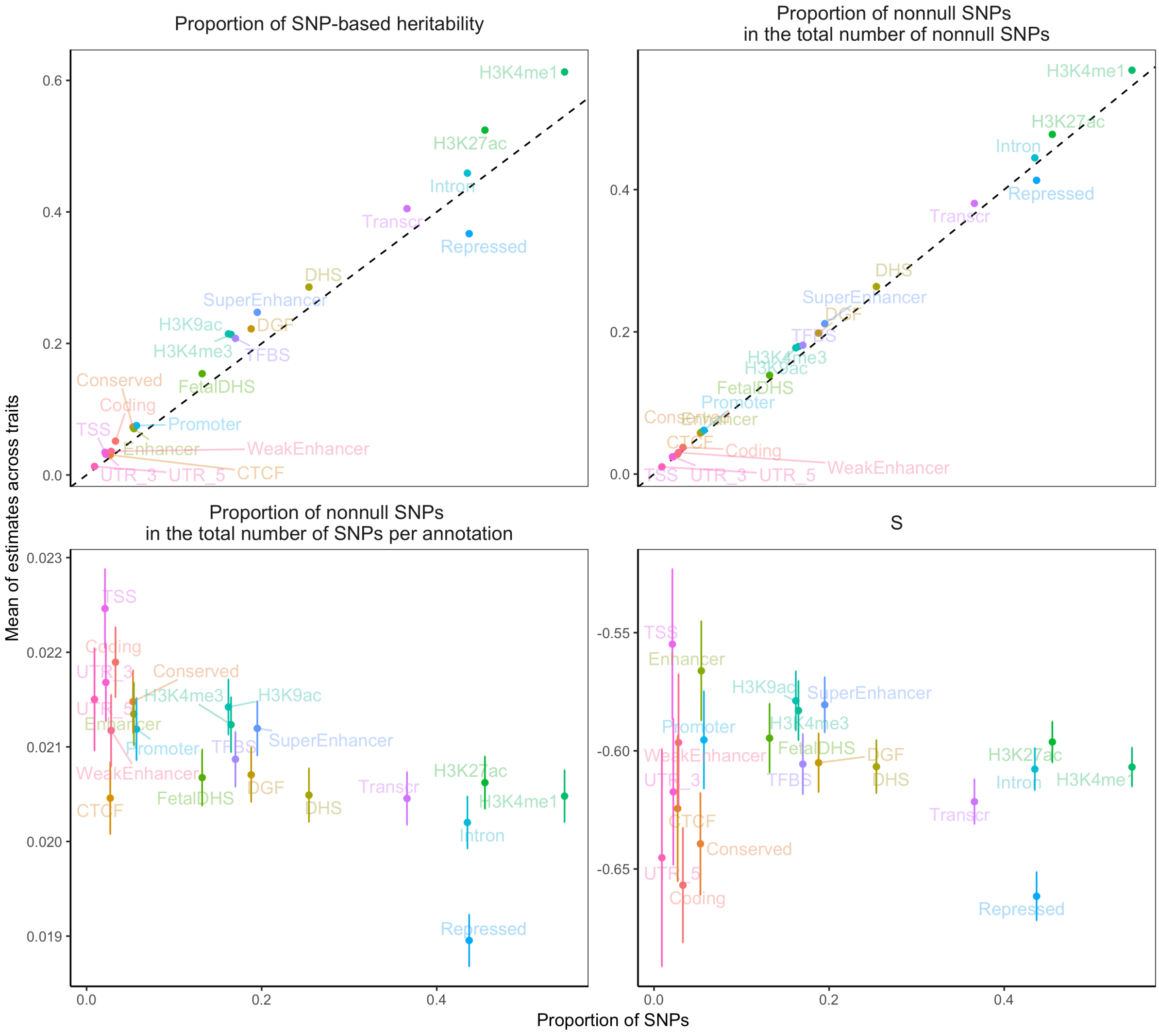


**Supplementary Figure 13** Estimation of the mean genetic architecture parameters across 35 UKB complex traits and diseases for the 21 functional annotation categories. The x-axis is the proportion of 1.1 million common HapMap3 SNPs that were allocated in each annotation. Each bar indicates the posterior standard error of the mean. Functional annotation categories are shown by text and colours. Dash line is the identity line.

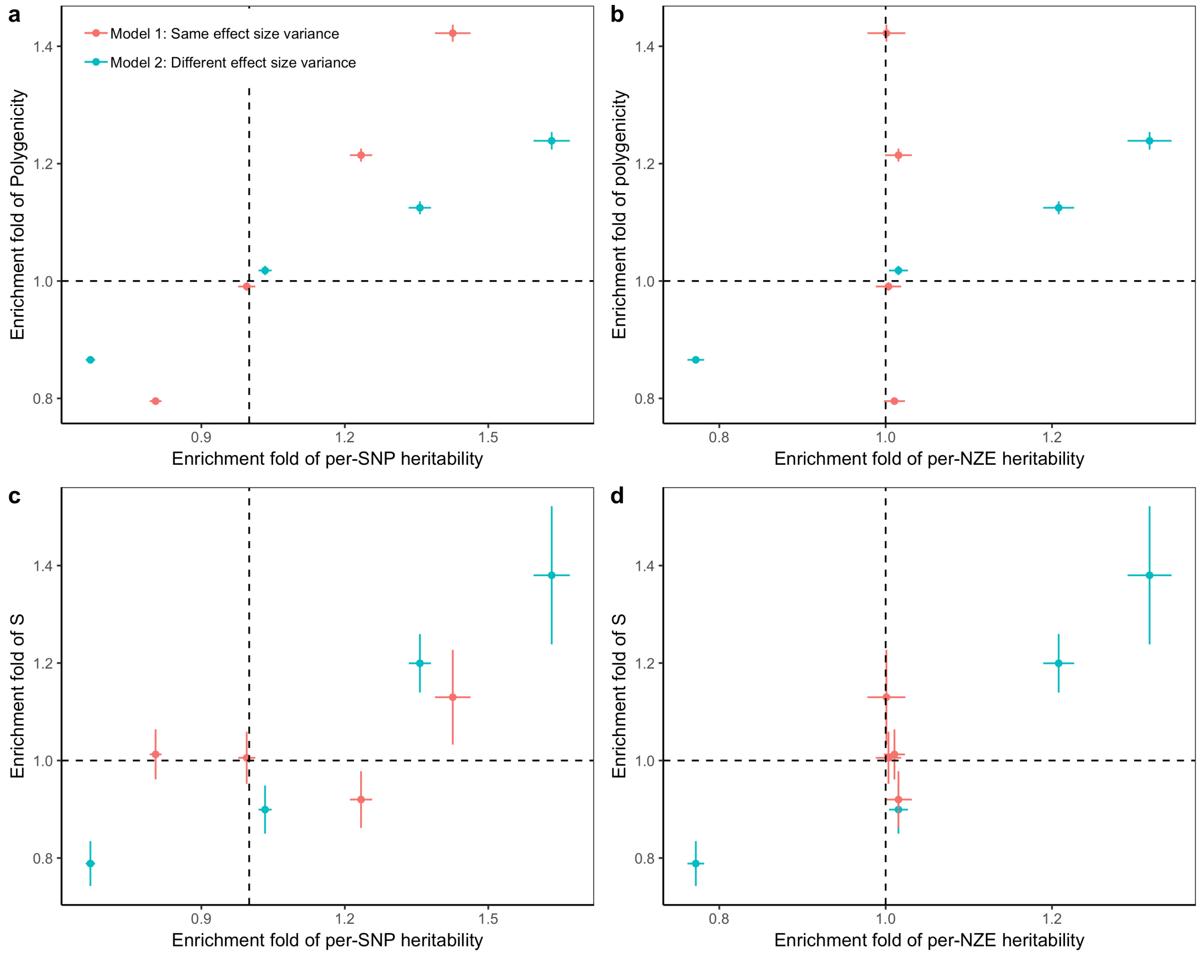


**Supplementary Figure 14** Estimation of parameter enrichment under different models of genetic architecture in forward simulations. In the simulation, a 10 Mb sequence consisted of four regions with sizes of 4, 3, 2 and 1 Mb, respectively. The proportion of trait mutations was 4%, 5%, 6% and 7% for the 4 regions. The trait heritability was set to be 0.05. A model of normalising stabilising selection was simulated on a population with a constant size of 10,000 for 10,000 generations. Two models were simulated for the distribution of causal effects, which followed a normal distribution with mean zero. In model 1, the causal effects have identical variance across regions. In model 2, the causal effects have variance of 1.4, 1.2, 1.0 and 0.8 for region 1 to 4, respectively. The results showed that the enrichment in per-SNP heritability was observed in both models (a), while per-NZE heritability enrichment was observed only in model 2 (b). Moreover, the enrichment of $\hat{S}$ is expected in the region with larger effect variance (c,d).

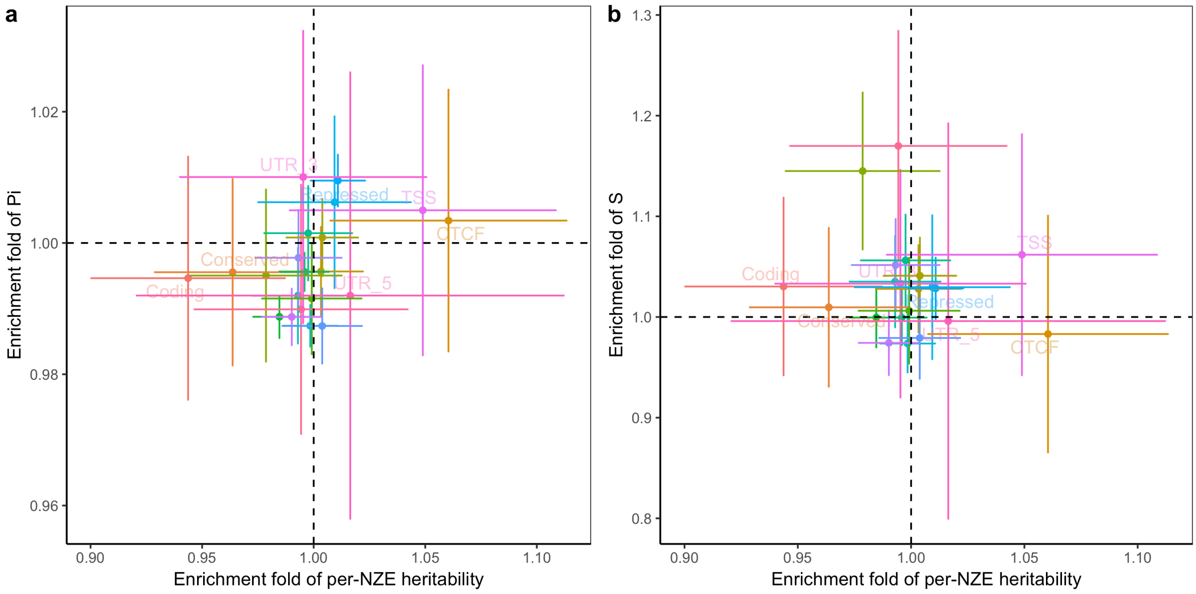


**Supplementary Figure 15** Estimation of enrichment in genetic architecture parameters for the 21 functional annotation categories under the null model. In this set of simulations, there were 5,000 causal variants randomly distributed in the genome regardless of the functional annotations. The heritability was set to be 0.5. The simulation was repeated 30 times. Each dot represents the mean across simulation replicates, and each bar is the standard error of the mean.

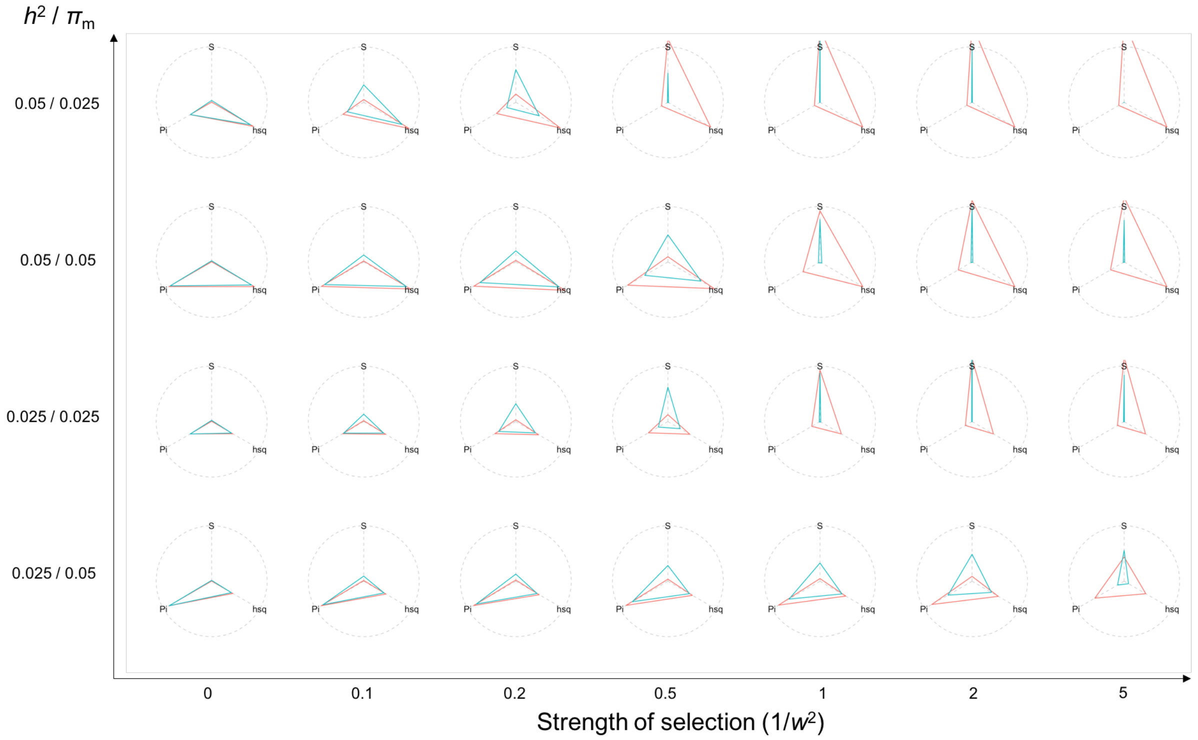


**Supplementary Figure 16** A map of genetic architecture patterns from forward simulations. Both x and y axes are input values in the simulation, where x-axis is the strength of selection denoted as one over the variance ($w^2$) of the phenotypes surrounding at the optimum fitness value (a classic model for stabilizing selection), and y-axis is the ratio of the heritability at all causal variants ($h^2$) over the proportion of mutational targets ($\pi_m$), which is proportional to the per-mutation heritability. Each circle consists of the values of SNP-based heritability, polygenicity and S, which are scaled by the maximum values across simulated traits such that the three parameters have the same scale from zero to one. The green (red) triangle indicates the computed values of the three genetic architecture parameters at the common (all) causal variants in the last generation of the forward simulation given the input values at x and y axes.

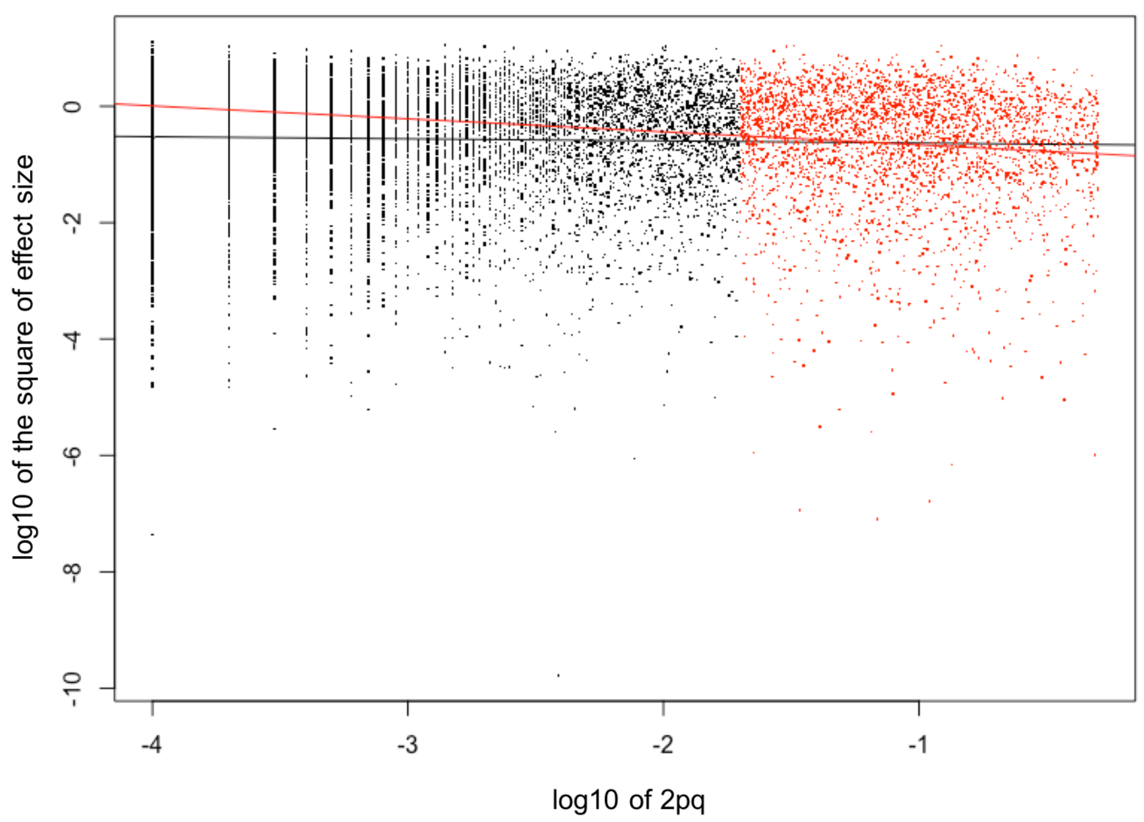


**Supplementary Figure 17** Joint distribution of the squared effect size and heterozygosity (2pq) for causal variants in the forward simulation in the presence of negative selection. Red colour shows the causal variants with MAF > 0.01. The line is the regression line for all causal variants (black) or common causal variants (red), which is an estimate of the *S* parameter.

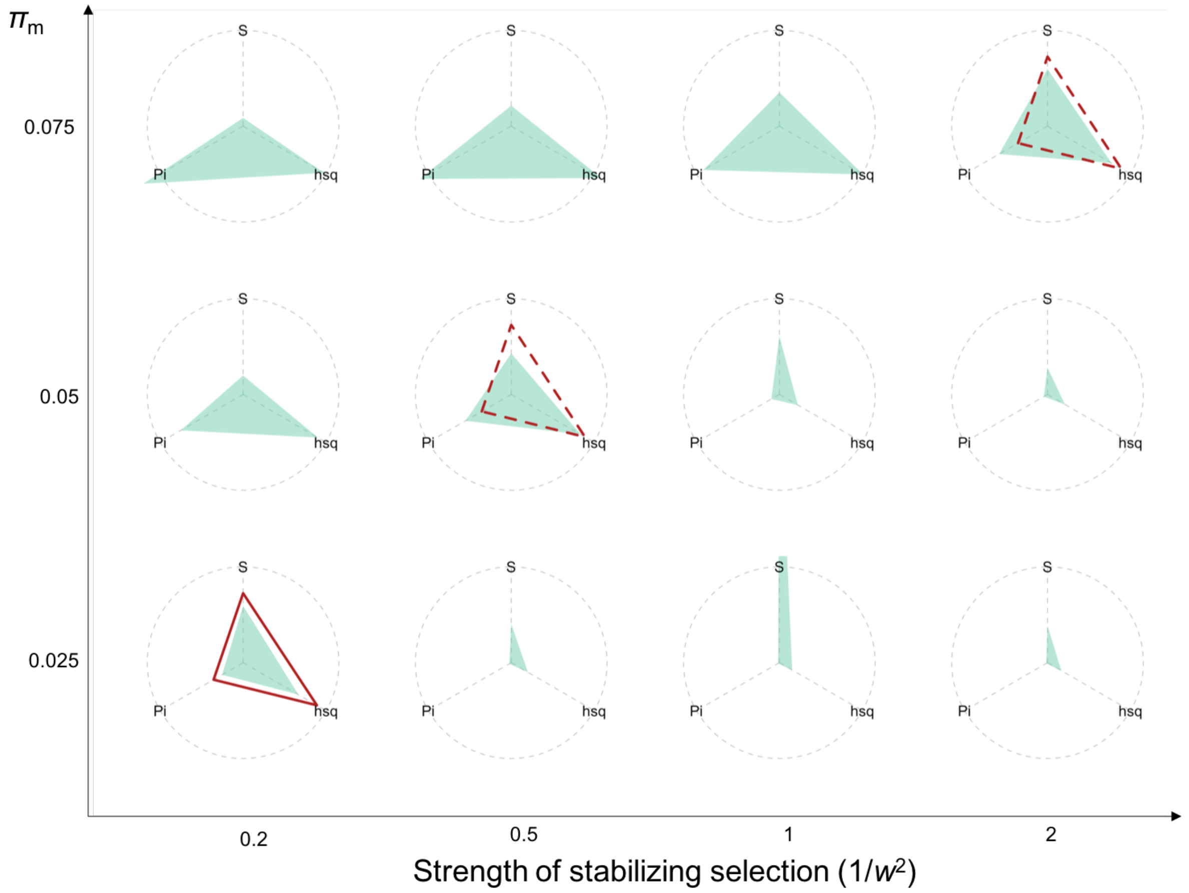


**Supplementary Figure 18** Projection of the genetic architecture for height onto a map of patterns from forward simulations incorporating both directional and stabilising selection. Both x and y axes are input values in the simulation, where x-axis is the strength of stabilising selection denoted as one over the variance ($w^2$) of the phenotypes surrounding at the optimum fitness value (a classic model for stabilizing selection), and y-axis is the proportion of mutational targets ($\pi_m$). The trait heritability for the 10 Mb sequence was set to 0.05 according to the projection result for height in Fig. 5. The directional selection was incorporated by assuming that individual fitness $f = \exp\left(-\frac{(y-\theta)^2}{2w^2}\right)$ where $y$ is the phenotype and $\theta = 0.2$ quantifies the strength of directional selection. The green shadow indicates the computed values for the three genetic architecture parameters at the common causal variants (MAF>0.01) in the last generation of the forward simulation given the input values at x and y axes. The solid (dashed) red triangle shows the best (suboptimal) projection of the estimated genetic architecture for height. Results suggest that in the presence of positive selection, genetic variants associated with height could appear to be under the same strength of negative selection as those with BMI ($1/w^2 = 2$ according to Fig. 4) when $\pi_m$ for height (0.075) is larger than that for BMI (0.05 in Fig. 4).

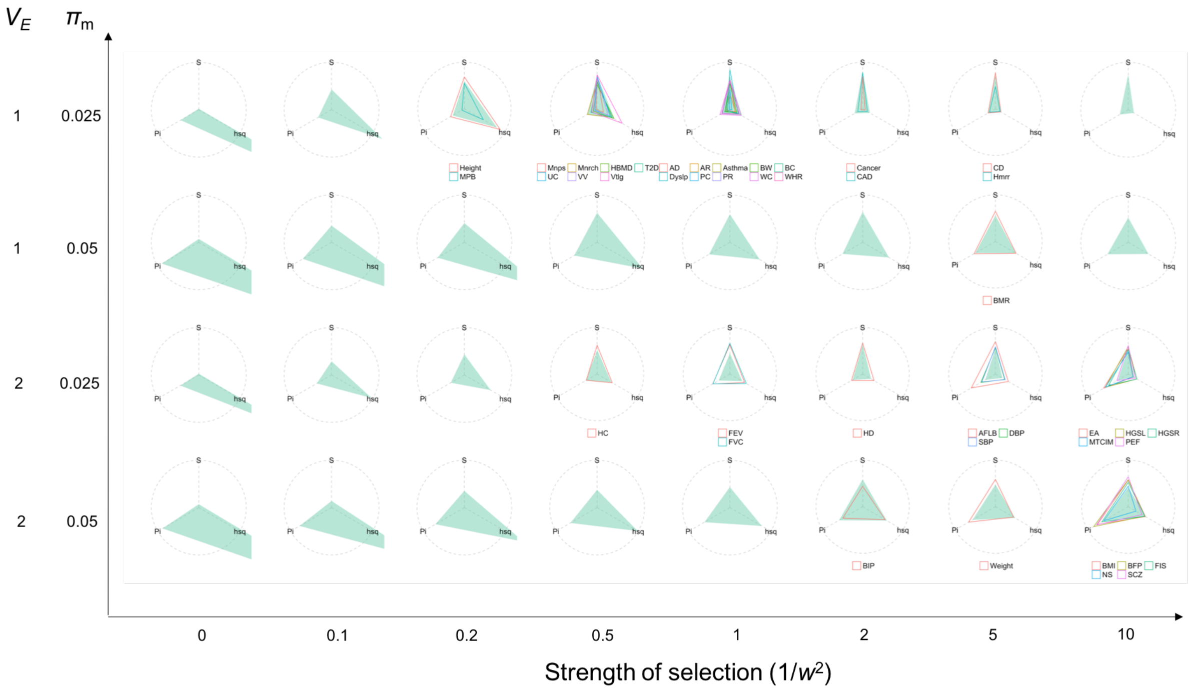


**Supplementary Figure 19** Projection of real trait genetic architecture onto the patterns observed from forward simulations with a demographic model and a constant environmental variance over generations. Both x and y axes are input values in the simulation, where x-axis is the strength of selection denoted as one over the variance ($w^2$) of the phenotypes surrounding at the optimum fitness value (a classic model for stabilizing selection), and y-axis is the ratio of the environmental variance ($V_E$) over the proportion of mutational targets ($\pi_m$), which is proportional to the per-mutation heritability. In the simulation, causal effects were sampled from $N(0, 0.01)$. Each circle consists of the values of SNP-based heritability, polygenicity and $S$, which are scaled by the maximum values across simulated or real traits such that the three parameters have the same scale from zero to one (note that the heritability in the simulation was scaled by 0.05 so that the results can be compared to those in Figure 4 on the same scale). The green shadow indicates the computed values for the three genetic architecture parameters at the common causal variants (MAF>0.01) in the last generation of the forward simulation given the input values at x and y axes. The hollow triangle shows the estimated genetic architecture for a UKB trait. The projection of UKB trait to the map from simulation was done by minimizing the sum of the difference in each of the three angles. The forward simulation was based on a 10 Mb sequence and repeated 30 times in each scenario. When there is no selection ($1/w^2 = 0$), as expected, the heritability tends to be close to one and $S = 0$.

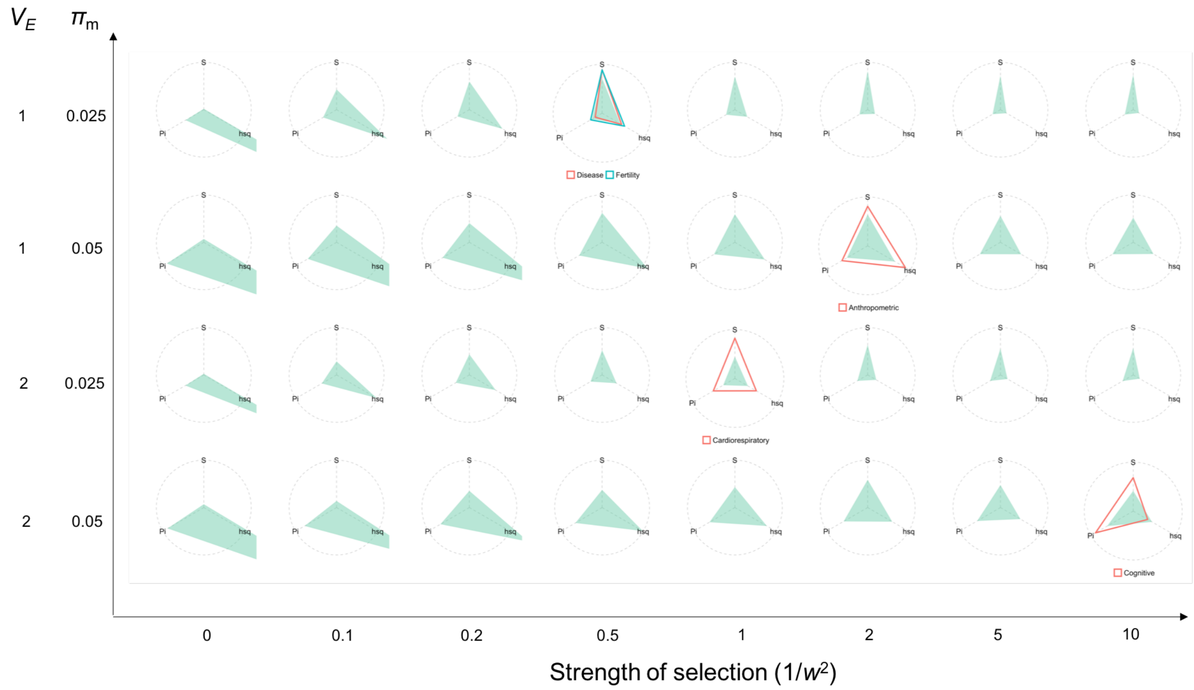


**Supplementary Figure 20** Projection of the observed genetic architecture averaged over each trait category onto the patterns observed from forward simulations with a demographic model and a constant environmental variance over generations. Both x and y axes are input values in the simulation, where x-axis is the strength of selection denoted as one over the variance ($w^2$) of the phenotypes surrounding at the optimum fitness value (a classic model for stabilizing selection), and y-axis is the ratio of the environmental variance ($V_E$) over the proportion of mutational targets ($\pi_m$), which is proportional to the per-mutation heritability. In the simulation, causal effects were sampled from $N(0, 0.01)$. Each circle consists of the values of SNP-based heritability, polygenicity and $S$, which are scaled by the maximum values across simulated or real traits such that the three parameters have the same scale from zero to one (note that the heritability in the simulation was scaled by 0.05 so that the results can be compared to those in Figure 5a on the same scale). The green shadow indicates the computed values for the three genetic architecture parameters at the common causal variants (MAF>0.01) in the last generation of the forward simulation given the input values at x and y axes. The hollow triangle shows the estimated genetic architecture for each trait category. The projection of the observed genetic architecture to the map from simulation was done by minimizing the sum of the difference in each of the three angles. The forward simulation was based on a 10 Mb sequence and repeated 30 times in each scenario. When there is no selection ($1/w^2 = 0$), as expected, the heritability tends to be close to one and $S = 0$.

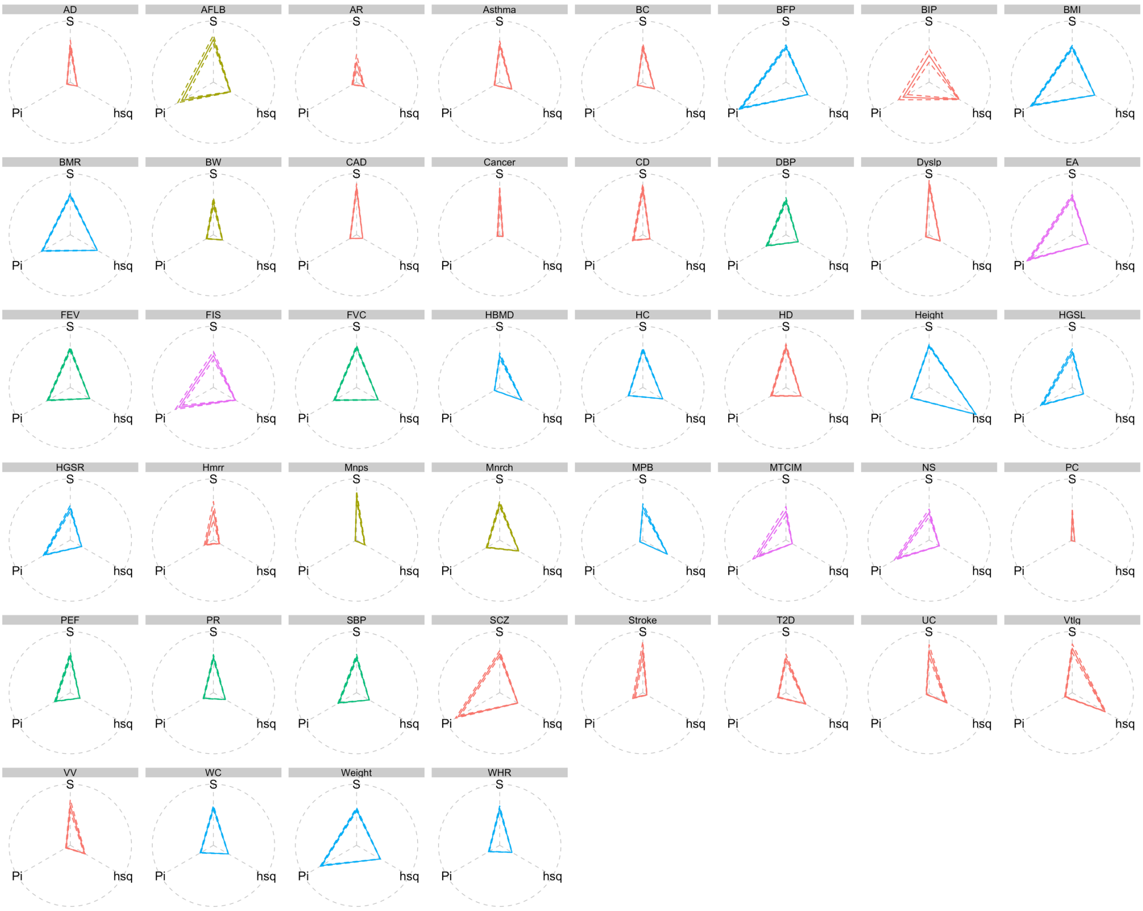


**Supplementary Figure 21** Genetic architecture parameter estimates for the 44 UKB traits (including diseases). The estimates of SNP-based heritability, polygenicity and *S* for each trait were scaled by the maximum values across traits and shown in a radar plot. Dash lines connect the standard errors of the estimates.
